# Introduction of regioselective bacterial heme oxygenases into *Arabidopsis hy1-1* supports the retrograde heme signaling hypothesis

**DOI:** 10.1101/2025.11.09.686882

**Authors:** Mihoko Takenoya, Takayuki Shimizu, Keita Miyake, Tatsuru Masuda

## Abstract

Heme is synthesized in the plastid and catabolized by heme oxygenase (HO). In *Arabidopsis*, HO1/HY1/GUN2 predominantly functions for heme catabolism. Previously, we demonstrated that introduction of either plastid-or cytosol-localized HO1 into *gun2* or *hy1-1* complemented the phenotypes including *genomes uncoupling* (*gun*), indicating the assembly of functional phytochromes, as well as supporting the heme signaling hypothesis.

To dissect the heme signaling and phytochrome assembly, in this study, we introduced either of the two types of regiospecific bacterial HOs that produce biliverdin IXα (BVIXα) or BVIXβ/δ into *hy1-1*. Introduction of BVIXα-producing HO complemented the long hypocotyl, low pigmentation, and *gun* phenotypes of *hy1-1*. Interestingly, the introduction of BVIXβ/δ-producing HO failed to complement the long hypocotyl and low pigmentation phenotypes, suggesting failure of functional phytochrome assembly. However, these lines restored the *gun* phenotype, thus supporting the heme signaling hypothesis.

Based on the levels of complementation of the *gun* phenotype, we found that the expression of photosynthesis-associated nuclear genes (*PhANGs*) can be separated into phytochrome-dependent and –independent groups. Our results demonstrate that heme functions as a retrograde mobile biogenic signal from plastids, passing through the cytosol, to regulate the expression of *PhANGs*, and this regulation is distinct in its dependency of phytochromes.

## Introduction

Heme serves as a cofactor for hemoproteins in various organelles, functioning in mitochondrial respiration and chloroplast photosynthetic electron transport chains, in detoxification of reactive oxygen species and xenobiotics, and in oxygen storage and transport (Layer et al., 2010). In addition, heme has been proposed to be a regulatory factor in transcriptional control and intracellular signaling in yeast and animals (Mense and Zhang, 2006; Tsiftsoglou et al., 2006).

Heme is synthesized through a biosynthetic pathway that begins with the universal precursor 5-aminolevulinic acid (ALA) (Beale, 1999) (Fig. 1). ALA is further metabolized through a series of enzymatic steps to form protoporphyrin IX, where the pathway is divided into heme and chlorophyll (Chl) branches (Tanaka et al., 2011). In the first step of the heme branch, ferrochelatase (FC) inserts Fe^2+^ into protoporphyrin IX to form heme (protoheme or heme *b*). Angiosperms have 2 *FC* genes, *FC1* and *FC2*, that exhibit distinct tissue– and development-dependent expression profiles (Tanaka et al., 2011). In mammals and yeast, FC is localized in the mitochondria, whereas in plant cells, both FCs are localized in the plastids (Masuda et al., 2003; Mochizuki et al., 2010). After the synthesis in plastids, heme is further catabolized by two enzymatic steps: the conversion of heme to biliverdin (BV) IXα by heme oxygenase (HO) (Weller et al., 1996; Weller et al., 1997) and the reduction of BVIXα to phytochromobilin (PΦB) by PΦB synthase (HY2) (Terry et al., 1995). Among the *Arabidopsis* 4 HO isoforms, HO1/HY1/GUN2 predominantly functions for heme catabolism. Other members, HO3 and HO4, are involved in HO1 family and have additional activities for BVIXα production with HO1 (Emborg et al., 2006; Gisk et al., 2010). On the other hand, HO2 cannot bind and convert heme and shows strong affinity to protoporphyrin IX. It is proposed that HO2 may be involved in the regulation of tetrapyrrole biosynthesis (Gisk et al., 2010). Thus, the HO1– and HY2-dependent pathway is predominant for holo-phytochrome (PHY) biosynthesis, since plant PHYs require thioether-linked PΦB prosthetic group as chromophore for light perception (Lagarias and Rapoport, 1980; Cornejo et al., 1992).

**Figure 1.**
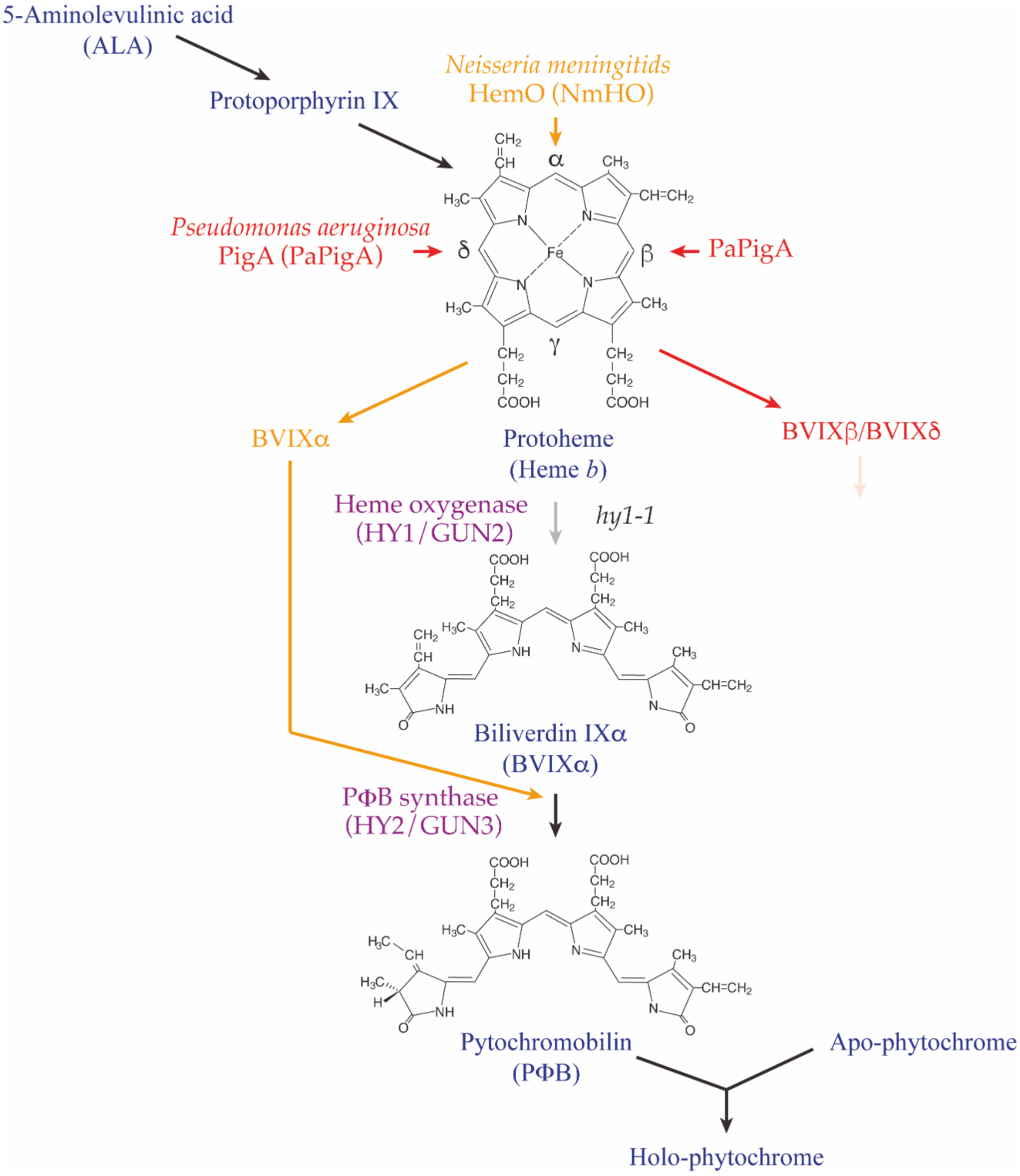
The heme branch pathway in plants consists of ferrochelatase (FC), heme oxygenase (HO), and phytochromobilin (PΦB) synthase (HY2). Synthesized PΦB binds to apo-phytochrome (PHY) to assemble functional holo-PHY. Among the 4 HO isoforms, HO1/HY1/GUN2 predominantly functions for heme catabolism. *hy1-1* and *hy2* are *HO1*– and *HY2*-deficient *Arabidopsis* mutants, respectively. *Neisseria meningitids* HemO (NmHO) cleaves the meso-α position of protoheme to produce biliverdin (BV) IXα (shown in orange), which may complement the *HO1*-deficiency of *hy1-1*. Meanwhile, *Pseudomonas aeruginosa* (*Pa*) *PigA-encoded* HO (PaHO) cleaves the meso-β or meso-δ position of heme, producing BVIXβ/δ (shown in red), which may not complement *hy1-1* while degrading heme.

Since HO1 and HY2 possess the N-terminal transit peptide and are ferredoxin-dependent reductases that obtain electrons from photosynthetic electron transport in chloroplasts (Muramoto et al., 1999; Kohchi et al., 2001; Muramoto et al., 2002), it is believed that PΦB synthesis occurs only in plastids (Terry et al., 1993; Kohchi et al., 2001). However, our previous study demonstrated that HO1 localization is altered in plastids or the cytosol due to regulation of the transcriptional start site (TSS) (Chen et al., 2024). Introduction of either plastid-or cytosol-localized HO1 into *HO1*-deficient mutants (*gun2* and *hy1-1*) resulted in recovery from the long hypocotyl and low pigmentation. Furthermore, high accumulation of PHY proteins in *gun2-1* was also restored in the transgenic lines, indicating the assembly of functional PHYs within these lines. The TSS-dependent cytosol and plastid HO1-dependent heme catabolism is regulated by light and development, and the cytosol pathway predominantly functions during early and skotomorphogenic development. In contrast, the plastid pathway is active in light-matured chloroplasts (Chen et al., 2024).

The nuclear genome encodes various components required for plastid structure and development. Thus, chloroplast biogenesis requires the coordinated expression of the plastid and nuclear genomes, with information exchanging between the nucleus and plastids. The latter is achieved through plastid-to-nucleus (retrograde) signaling pathways, in which plastids signal to regulate various physiological processes, such as the expression of photosynthesis-associated nuclear genes (*PhANGs*), depending on their developmental and functional states. Although multiple retrograde signaling pathways have been proposed, “biogenic control” signals are thought to act during the initial stage of chloroplast development (Pogson et al., 2008). Genetic and biochemical analyses of the biogenic retrograde pathway suggest a significant role for heme in retrograde signaling (Shimizu and Masuda, 2021). In *Arabidopsis*, mutations affecting chloroplast function or treatments with inhibitors such as norflurazon (NF) or lincomycin (Lin) result in the intense repression of many *PhANGs*. The identification of *genomes uncoupled* (*gun*) mutants, in which the expression of *PhANGs* is maintained following chloroplast damage induced by inhibitor treatment, suggests the involvement of tetrapyrroles in retrograde signaling (Susek et al., 1993). Among the original five *gun* mutants described, *gun2* and *gun3* lack a functional HO1 and HY2, both of which are involved in the heme branch (Tanaka et al., 2011).

Meanwhile, *gun4* and *gun5* are mutants of the regulator and the catalytic CHLH subunit of Mg-chelatase, respectively (Mochizuki et al., 2001). Subsequently, the identification of a dominant *gun6* mutant with increased FC1 activity has restored *PhANGs* expression, even when chloroplast development is blocked (Woodson et al., 2011; Page et al., 2020). In contrast, the overexpression of *FC2* was not effective for the restoration. These data suggest that increased flux through the FC1-producing heme may act as a signaling molecule that positively controls *PhANGs* as a retrograde signal in *Arabidopsis* (Shimizu and Masuda, 2021). Subsequently, we demonstrated that GUN1, which encodes a plastid-localized pentatricopeptide repeat protein functioning as a master switch that integrates multiple retrograde signaling pathways (Koussevitzky et al., 2007), can directly bind heme and modulate tetrapyrrole biosynthesis (Shimizu et al., 2019). It is proposed that GUN1 regulates FC1-derived heme transfer from plastids at the initial stage of chloroplast development (Shimizu and Masuda, 2021). Our previous study also showed that the overexpression of *HO1* in the cytosol or plastids under the *gun2-1* background complemented the *gun* phenotype, confirming that heme degradation in the cytosol or plastids likely canceled the positive effect on *PhANGs* expression (Chen et al., 2024).

However, to demonstrate the heme signaling hypothesis as a biogenic retrograde signaling, a possibility for crosstalk with light signaling in *PhANGs* expression (Larkin and Ruckle, 2008; Larkin, 2014) needs to be evaluated. The perception of light signals by the PHYs and cryptochrome (CRY) family of photoreceptors has a crucial influence on various aspects of plant growth and development. The *CRY1*-deficient mutant (*cry1*) was identified to exhibit a *gun* phenotype (Ruckle et al., 2007). Concerning the involvement of PHYs in the retrograde *gun* signaling, *gun4* and *gun5* mutants deficient in a regulator and catalytic subunit of Mg-chelatase may produce functional PHY (Mochizuki et al., 2001). In addition, *phyA* and *phyB* mutants did not exhibit the *gun* phenotype (Ruckle et al., 2007). However, it is proposed that PHYB becomes a negative regulator in a *gun1* background (Ruckle et al., 2007). Since the CRY1 and PHYs interaction predominantly affects the *PhANGs* and photomorphogenesis, it is proposed that the crosstalk between plastid and light signaling pathways that affect *PhANGs* expression when chloroplast biogenesis is blocked may involve one or more PHYs (Ruckle et al., 2007).

To clarify this possibility, we introduced regiospecific HOs into the *Arabidopsis hy1-1* background in this study. Many pathogenic bacteria possess specific heme uptake systems that utilize heme-iron as nutrients. Bacterial heme assimilation involves the uptake of intact heme into the cell (Wandersman and Stojiljkovic, 2000). The oxidative cleavage of incorporated heme by HO releases iron, which can be utilized by the cell as an iron source. Although most bacterial HOs produce BVIXα as a product, the *Pseudomonas aeruginosa* (*Pa*) *PigA-encoded* HO (*PaHO*) was found to produce BVIXβ and BVIXδ as major and minor products, respectively, showing PaHO is an exceptional HO with a novel regiospecificity (Ratliff et al., 2001) (Fig. 1). The regiospecificity of PaHO is caused by the unusual seating of the heme in the enzyme, with a rotation in-plane of ∼110° compared to BVIXα-producing HO (Caignan et al., 2002). Since BVIXβ/δ may not be used as substrates for PΦB-producing HY2, we hypothesize that if *PaHO* is introduced into *hy1-1*, functional holo-PHY may not be assembled, as heme is catabolized to BVIXβ/δ. This may allow us to analyze how the functional holo-PHY assembly is involved in the biogenic retrograde signaling. In this study, we compared the effects of introducing BVIXα– and BVIXβ/δ-producing HO in the *HO1*-deficient mutant *hy1-1*. For this purpose, we utilized *Neisseria meningitidis* HO (NmHO) (Zhu et al., 2000) as a BVIXα-producing HO and PaHO as a BVIXβ/δ-producing HO (Fig. 1).

## Materials and Methods

### Plant materials and growth conditions

*Arabidopsis* (*A. thaliana*) Ler and mutants *hy1-1* were used in this research. *Arabidopsis* seeds were sterilized and sown on 2/3 Murashige and Skoog (MS) medium. Subsequently, they underwent a 1-day cold treatment before being transferred to 22°C and exposed to white light (40 μmol photons m^−2^ s^−1^) for 4 days to induce germination. The LA-105 Light Analyzer (NK system, Japan) was used to measure light intensities. NF (Fujifilm, Japan) was added directly to the MS medium at a final concentration of 5 μM. For the timing of activation of the biogenic retrograde signaling examination, *Arabidopsis* Ler seeds were sterilized and exposed to the same white light conditions as in liquid MS medium. Liquid cultures are exposed to the same light intensity with shaking (∼80 rpm). NF was added at the indicated time of culture.

### Docking simulation

Each biliverdin isomer (BVIXα: CID 5280353, BVIXβ: CID 126456549, and BVIXδ: CID 126456551) was prepared using the corresponding SDF structures downloaded from PubChem. The ligand structures were converted into AutoDock Vina–compatible PDBQT format using the Meeko package (v0.6.1). The receptor structure used for docking was tomato HY2 (PDB ID: 6KME) (Sugishima et al., 2020). Chain A was extracted with the ProDy package (Bakan et al., 2011; Zhang et al., 2021) and hydrogen atoms were added using reduce2.py (mmtbx module) with reference to the CCP4 monomer library (geostd). Water molecules located within 3 Å of the native ligand (BVIXα) were retained. The processed receptor was then converted into PDBQT format using Meeko’s mk_prepare_receptor.py for subsequent docking. Docking simulations were performed using AutoDock Vina (ver. 1.2.6) (Trott and Olson, 2010; Eberhardt et al., 2021) Two docking modes were examined: a box-restricted mode, in which the search box (20 Å per side) was centered at the centroid of the native ligand (BVIXα), and a blind docking mode, in which the search region was extended to the entire protein structure. The docking parameters were set as follows: exhaustiveness = 54, num_modes = 50, energy_range = 4, and seed = 10.

### Vector construction and plant transformation

Amino acid sequences of NmHO and PaHO were obtained from Ratliff et al. (2001). Based on these sequences, codon-optimized nucleotide sequences for *Arabidopsis* were generated by VectorBuilder (https://en.vectorbuilder.com/) (Supplemental Fig. 1). These sequences were artificially synthesized and subcloned into the pEFk vector by Fasmac (https://www.bio.fasmac.co.jp/), which were directly transformed into *Escherichia coli* (*E. coli*) BL21(DE3) for recombinant protein expression. To construct transgenic *Arabidopsis*, the coding sequences were inserted into the pENTR/D-TOPO vector (Invitrogen, USA) according to the manufacturer’s instructions, yielding pENTR/D-TOPO-NmHO and pENTR/D-TOPO-PaHO. The transit peptide sequence of the *Arabidopsis* small subunit of ribulose-bisphosphate carboxylase/oxygenase (RBCS) was inserted before the initiation codon of bacterial HOs by Gibson assembly® cloning (New England Biolabs, USA), producing pENTR/D-TOPO-cNmHO and pENTR/D-TOPO-cPaHO (Supplementary Fig. 2). These gateway entry clones were subjected to an LR recombination reaction (Invitrogen) with pGWB5 to construct the corresponding expression vectors, pGWB5/p35S::NmHO-GFP:nopaline synthase terminator (tNOS), pGWB5/p35S::PaHO-GFP:tNOS, pGWB5/p35S::cNmHO-GFP:tNOS, and pGWB5/p35S::cPaHO-GFP:tNOS. The nucleotide sequences of the inserted genes in all the plasmids described above were confirmed by sequencing analysis. PCR primers are listed in Supplementary Table S1.

The obtained destination vectors were transformed into *Agrobacterium tumefaciens* GV3101 using the freeze–thaw method (Weigel and Glazebrook, 2006). The GV3101 strain containing target vectors was transformed into *hy1-1* via *Agrobacterium*-mediated floral dip transformation (Zhang et al., 2006). T0 to T3 generations were screened on the MS medium containing 100 μg/ml cefotaxime and 50 μg/ml kanamycin.

### Protein expression and purification

The *E. coli* BL21(DE3) strain carrying pEFk vectors harboring codon-optimized *NmHO* and *PaHO* was grown in 150 ml of LB medium containing 50 µg/ml kanamycin to mid-log phase. Protein expression was induced by the addition of isopropyl-1-thiol-(D)-galactopyranoside (IPTG) to a final concentration of 1 mM. Protein expression was induced for 5 h at 30°C, and cells were harvested by centrifugation. Cells were disrupted by ultrasonication in buffer A (20 mM Tris-HCl, pH 7.5, 20 mM imidazole, and 500 mM NaCl) and then centrifuged at 15,000xg for 30 min. The supernatants were harvested and purified using a Ni-NTA (Qiagen, Germany) column by gravity flow. The target proteins were eluted with buffer B (20 mM Tris-HCl, pH 7.5, 500 mM imidazole, and 500 mM NaCl). The eluent was then dialyzed against buffer C (20 mM Tris-HCl, pH 7.5).

### HO assay

The regiospecific bacterial HO activity was assayed in the presence of ascorbate as an exogenous reductant (Zhu et al., 2000; Ratliff et al., 2001). Hemin was added to purified bacterial HO to achieve a final 2:1 heme-protein ratio, forming the heme-HO complex. To a 1 ml reaction mixture, 5 mM ascorbic acid was added directly to the HO-heme complex (10 µM) in 20 mM Tris-HCl buffer (pH 7.5) to initiate the reaction. The absorption spectral changes between 300 and 750 nm were recorded over a 30 min period. The reaction was monitored with a V-730 spectrophotometer (JASCO, Japan). The absorption spectra of the final products were recorded after acidification with glacial acetic acid (200 µl) and 3 M HCl (200 µl). The acidified products were further analyzed by HPLC after extraction and methylation (Zhu et al., 2000). The products were extracted with 1 ml of chloroform, and the organic layer was washed 3 times with 1 ml of distilled water. The chloroform layer was removed in a vacuum concentrator. The resultant residue was dissolved in 1 ml of 4% sulfuric acid in methanol and esterified for 12 h at room temperature. The esters were diluted with 4 ml of distilled water and extracted into 1 ml of chloroform. The organic layer was washed further with distilled water and again removed in a vacuum concentrator. The residue was dissolved in HPLC solvent before HPLC analysis. The samples were analyzed on reverse-phase HPLC on an ODS-C18 (GL Science, Japan) column (4.6 x 250 mm) eluted with methanol:H_2_O (85:15) at a flow rate of 0.8 ml/min. The elution was monitored with a photodiode-array detector (JASCO, Japan). Peaks of BV were determined in the eluted order (Zhu et al., 2000; Ratliff et al., 2001): BVIXα (7.1 min), BVIXδ (7.9 min), and BVIXβ (8.2 min).

### RNA extraction and RT-qPCR analysis

The seedlings were harvested and immediately frozen in liquid nitrogen. Total RNA was extracted using a RNeasy Plant Mini Kit (Qiagen, Germany), including treatment with an RT Grade DNase set (Nippon Gene, Japan). For RT-qPCR analysis, 500 ng of total RNA was reverse transcribed into cDNA using PrimeScript RT Master Mix (Takara). RT-qPCR was performed using the corresponding primers listed in Supplementary Table S1, with THUNDERBIRD SYBR qPCR Mix (Toyobo, Japan) on a QuantStudio 1 Real-Time PCR System (Thermo Fisher, USA). For the timing of activation of the biogenic retrograde signaling examination, 70.0 ng total RNA was reverse transcribed into cDNA.

### Protoplast isolation and fluorescence microscopy observation

Protoplasts were isolated from 3-week-old fully expanded leaves (Lung et al., 2015). GFP and Chl fluorescence signals were detected using an FV3000 confocal laser-scanning microscope (Olympus, Japan). The excitation wavelength was 488 nm, and the emission wavelengths were 500-550 nm for GFP and 662-691 nm for Chl.

### Phenotypic analysis

Photographs of 4-day-old plants were taken after they were transferred to 1.0% (w/v) agarose plates. Hypocotyl lengths were determined by analyzing captured photographs with NIH ImageJ. 4-day-old seedlings were harvested and immersed in 1 ml of 80% (v/v) acetone overnight to extract Chls and carotenoids (Lichtenthaler, 1987). Debris was removed by centrifugation at 18,000 × g for 5 min. The absorbance was measured with a V-730 spectrophotometer (JASCO, Japan) at 663, 647, and 470 nm. The Chl *a* and *b* concentrations were calculated as described previously (Chen et al., 2024). Heme was extracted and measured from the samples according to the published protocol (Espinas et al., 2012).

### Western blot

4-day-old plants were harvested and frozen immediately in liquid nitrogen. After homogenization, total proteins were extracted in Laemmli buffer. Protein concentrations were determined by an RC DC Protein Assay (Bio-Rad). For SDS-PAGE, 10 μg of protein was separated on a 10% (w/v) polyacrylamide gel and subsequently immunoblotted with an anti-GFP (MBL, Japan) or anti-PHYA (PhytoAB, USA) antibody.

### Statistical analysis

All measurements were done with more than three replications. Statistical analysis was performed using R and RStudio. To perform multiple comparisons, the TukeyHSD test was applied after the one-way ANOVA. Different letters in the figures indicate significant differences in P<0.05.

## Results

### Docking simulation for HY2 and BV

First, we evaluated whether BVIXβ or BVIXδ becomes a potential substrate for PΦB-producing HY2. For this purpose, we performed a computer-based simulation to estimate the required energy for binding substrate pocket of HY2 and BVs based on the determined structure of BVIXα-binding HY2 of tomato (Sugishima et al., 2020). Asp123 and Asp263 of HY2 are suggested to mediate the reduction of BV via water molecules (Fig. 2a). To examine the potential influence of water molecules on substrate binding, we performed docking simulations while retaining water molecules located within 3 Å of the native ligand (BVIXα).

**Figure 2.**
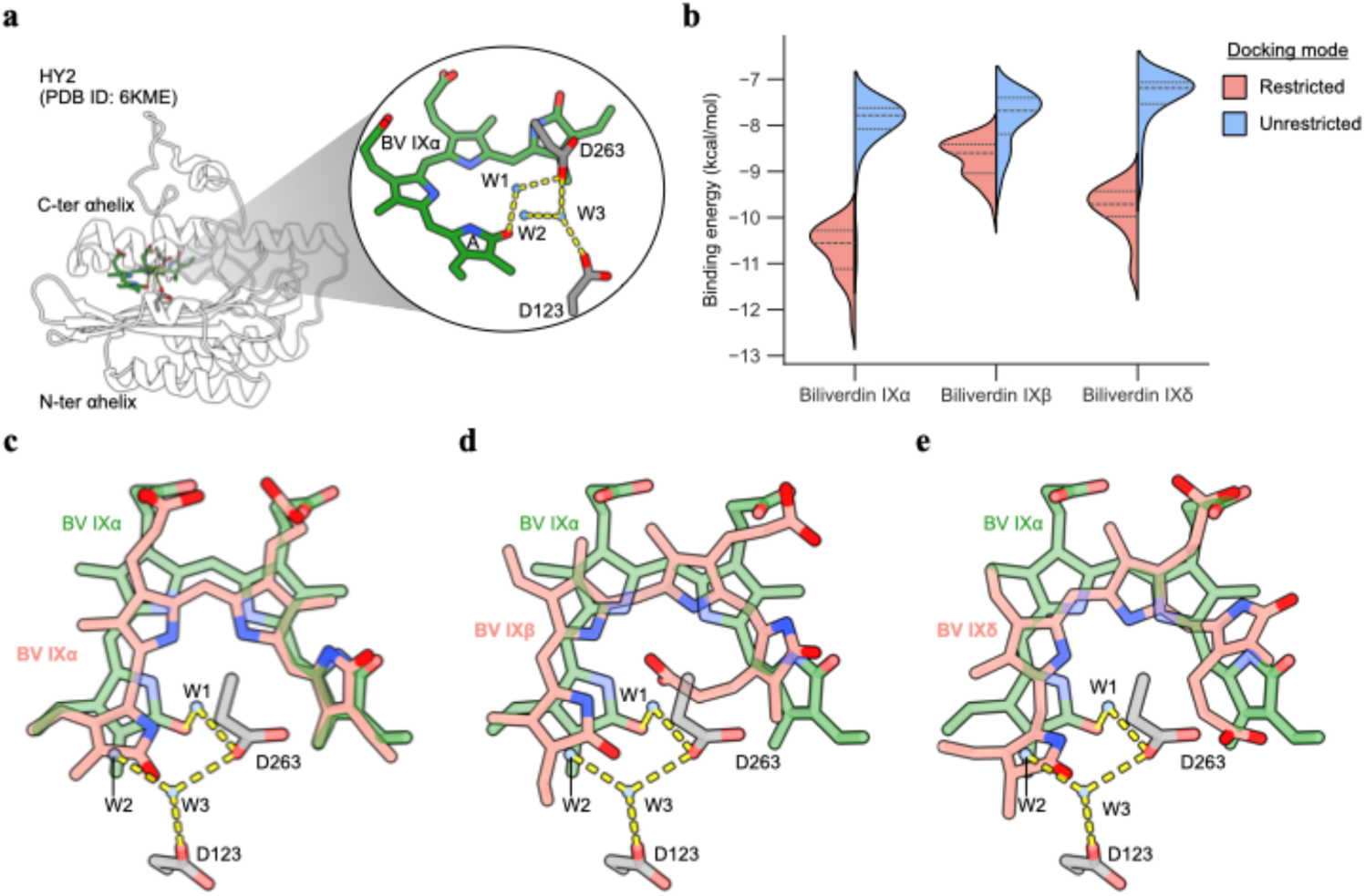
Docking simulation of HY2 with biliverdin isomers. (a) Overall structure of tomato HY2 (PDB ID: 6KME) showing Asp123 and Asp263, which are suggested to mediate biliverdin reduction through water molecules (W1-3). (c-e) Docking models of HY2 complexed with the lowest-energy conformations of BVIXα (c), BVIXβ (d), and BVIXδ (e) obtained under the restricted condition.

For all BV isomers, the binding energies were lower when docking was restricted to the substrate-binding pocket compared with the unrestricted mode (Fig. 2b). Among them, BVIXα showed the lowest binding energy under the restricted condition, indicating a more favorable interaction with HY2. The docking pose of BVIXα closely resembled that of the native ligand, although it was positioned slightly deeper into the pocket (Fig. 2c). In contrast, BVIXβ was accommodated with its carboxyl group located near the center of the binding pocket (Fig. 2d). In both BVIXβ and BVIXδ, the A– and D-rings were positioned close to Asp123, Asp263, and the intervening water molecules that are implicated in the reduction reaction (Fig. 2d, e). As shown in Supplement Fig. 1, BVIXβ and BVIXδ appeared not to fit well into the substrate pocket, whereas BVIXα was better accommodated within the binding pocket. Collectively, these results indicate that HY2 preferentially recognizes BVIXα, and that BVIXβ and BVIXδ are unlikely to serve as effective substrates.

### Enzyme activities of NmHO and PaHO

To introduce regiospecific two bacterial HO genes (*NmHO* and *PaHO*) into *Arabidopsis hy1-1*, we designed and synthesized codon-optimized artificial genes with identical amino acid sequences to the bacterial HOs (Supplemental Fig. 2). To confirm the regiospecific HO activities of NmHO and PaHO, we introduced them into an expression vector, expressed the proteins in *E. coli*, and purified them to homogeneity. Purified proteins were separated by SDS-PAGE with apparent bands of 26 kDa and 23 kDa for NmHO and PaHO, respectively (Fig. 3a). After mixing with hemin, the heme-HO complex was converted to BV in the presence of ascorbate as an exogenous reductant (Zhu et al., 2000). Both proteins showed heme-degrading activities (Fig. 3b, c). PaHO exhibited higher activity than that of NmHO, as heme degradation was almost completed within 10 min of incubation (Fig. 3c). In comparison, the heme peak (405 nm) remained after 20 min of incubation in NmHO (Fig. 3b). The absorption spectra of the final products showed peak maxima at 380 nm and 680 nm, indicating the presence of iron-free BV (Fig. 3d). Following the reaction, we analyzed the products by HPLC after extraction and methylation. HPLC analysis of the products of NmHO yielded a major peak with retention times identical to those of BVIXα (Fig. 3e). In contrast, the products of PaHO exhibited both minor and major peaks with longer retention times, which were previously identified as BVIXβ and BVIXδ, respectively (Ratliff et al., 2001) (Fig. 3e). In addition, only a minor peak of BVIXα was detected in the products. These results confirm that NmHO and PaHO are regiospecific HO-producing BVIXα and BVIXβ/δ, respectively.

**Figure 3.**
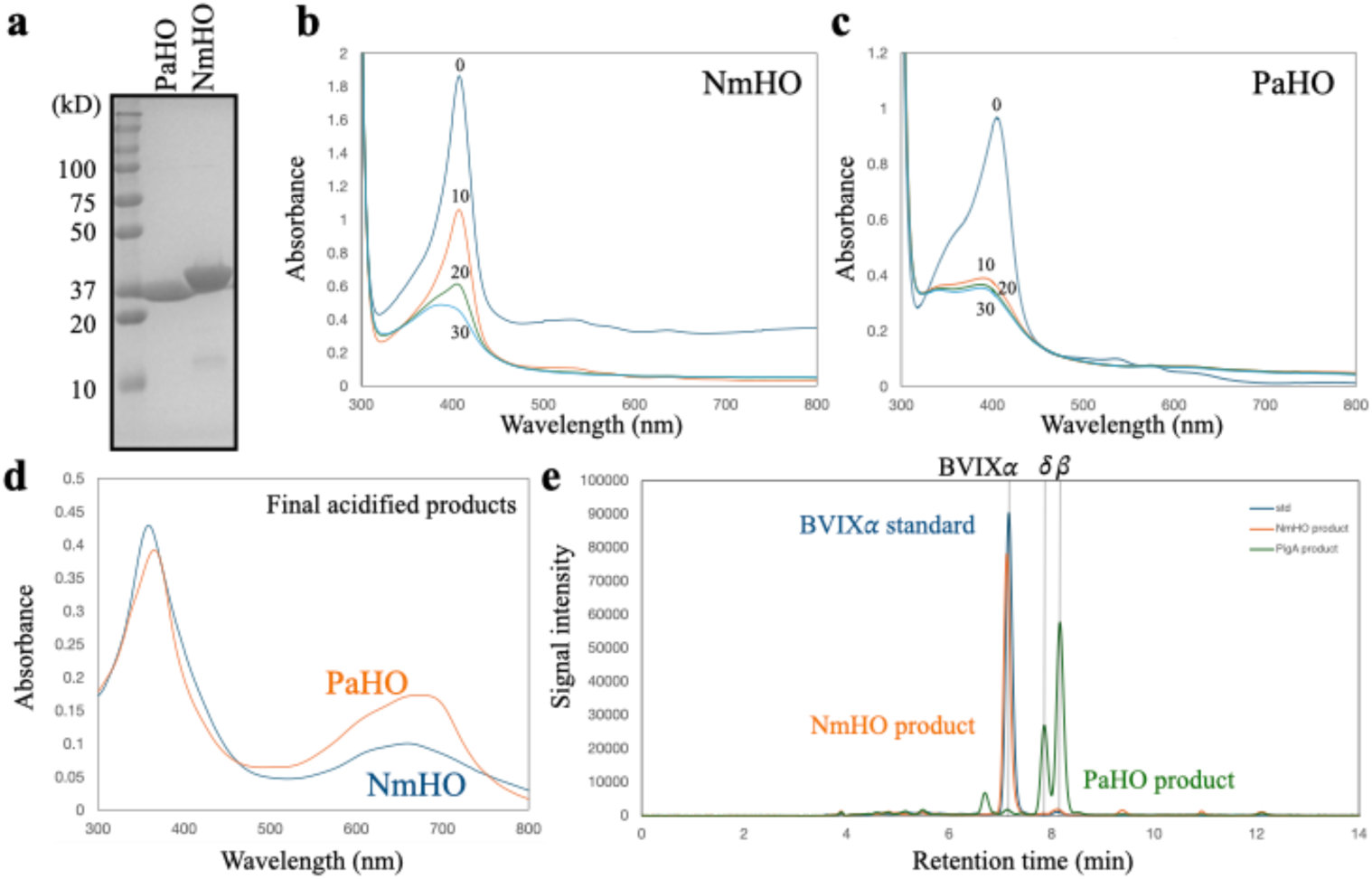
(a) CBB-stained gel of SDS-PAGE separated on purified NmHO and PaHO recombinant proteins. HO activity measurement of NmHO (b) and PaHO (c) by measuring the disappearance of heme absorption spectra every 10 min after reaction. Ascorbate was added as an exogenous reductant in this assay. (d) Absorption spectra of final BV products after acidification. (e) HPLC chromatogram of BV products. Elution profiles of BVIXα standard (blue), NmHO product (orange), and PaHO product (green) were depicted. Peaks of BV were determined in the eluted order (Zhu et al., 2000; Ratliff et al., 2001): BVIXα (7.1 min), BVIXδ (7.9 min), and BVIXβ (8.2 min).

### Construction of bacterial *HO* introduced *hy1-1* transgenic lines

To characterize the function of regiospecific HOs, we introduced them into the *Arabidopsis hy1-1*background. According to our previous study, which revealed the significance of cytosolic HO activity (Chen et al., 2024), we constructed *NmHO* and *PaHO* transgenic lines, either expressing them in plastids by adding an N-terminal RBCS transit peptide or in the cytosol without a transit peptide (Fig. 4a). As a result, we generated four transgenic lines expressing in plastids, *p35S::cNmHO-GFP/hy1-1* and *p35S::cPaHO-GFP/hy1-1* (hereafter referred to as the *cNmHO/hy1-1* and *cPaHO/hy1-1*), and cytosol, *p35S::NmHO-GFP/hy1-1* and *p35S::PaHO-GFP/hy1-1* (hereafter referred to as the *NmHO/hy1-1* and *PaHO/hy1-1*) (Fig. 4a).

**Figure 4.**
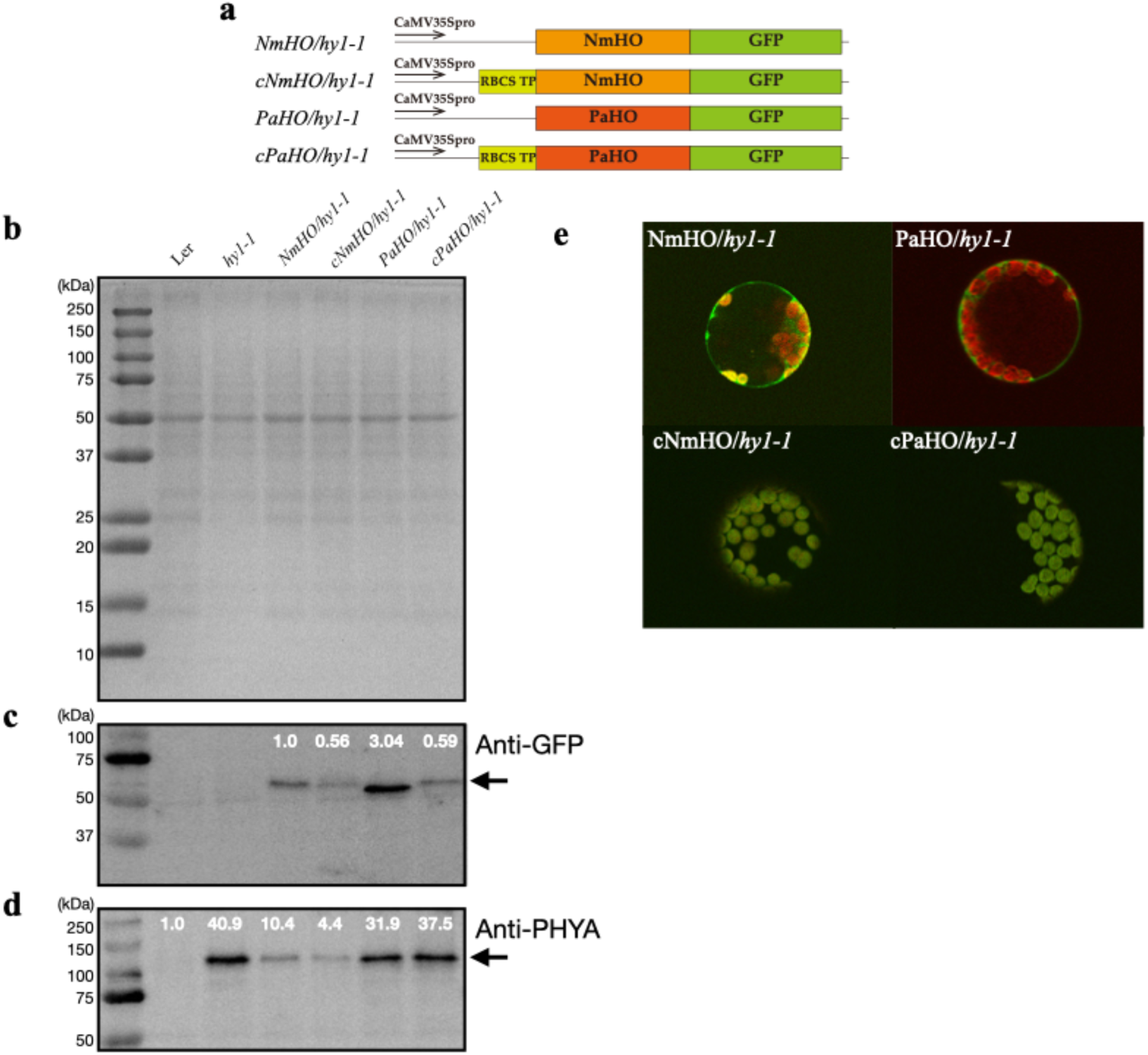
(a) Schematic diagram of transgenic lines. (b) CBB-stained SDS-PAGE gel of extracted total proteins from *Arabidopsis* seedlings. Western blot analysis with anti-GFP antibody (c) and anti-PHYA antibody (d). The white number above each band represents band intensity, quantified in ImageJ and normalized to RbcL, the large subunit of RuBisCO. Relative intensities to *NmHO/hy1-1* for GFP (c) and to wild-type Ler for PHYA (d) are indicated. (e) Merged images of EGFP and Chl fluorescence in protoplasts obtained from each transgenic line.

From 4-day-old light-grown wild-type, *hy1-1*, and transgenic lines, total proteins were extracted and separated by SDS-PAGE (Fig. 4b). We then performed Western blot analysis using anti-GFP and anti-PHYA antibodies. The expression levels of bacterial HOs were measured by Western blot using an anti-GFP antibody (Fig. 4c). The expression of transgenes was confirmed by the detection of a GFP signal only in transgenic lines. GFP signals were higher in the cytosolic lines (*NmHO/hy1-1* and *PaHO/hy1-1*) than in the respective plastid lines (*cNmHO/hy1-1* and *cPaHO/hy1-1*) (Fig. 4c). Western blot analysis of PHYA confirmed high accumulation of PHYA protein in the *hy1-1*, compared to the wild-type (Fig. 4d), which is consistent with the fact that the *HO1*-deficient mutant contains dysfunctional PHY apoproteins (Parks et al., 1989; Chen et al., 2024). The PHYA signal was low in *NmHO/hy1-1* and *cNmHO/hy1-1*, however the band intensities were still higher than that of wild-type. On the other hand, *PaHO/hy1-1* and *cPaHO/hy1-1* showed comparable signal intensities to that of *hy1-1* (Fig. 4d). This result confirms our hypothesis that, unlike BVIXα-producing NmHO, BVIXβ/δ-producing PaHO cannot assemble the functional PHY. Then, we confirmed the localization of *NmHO* and *PaHO* products by fluorescence microscopy imaging using protoplasts prepared from well-expanded leaves of the transgenic lines. As shown in Fig. 4e, GFP signals were detected outside of chloroplasts in *NmHO/hy1-1* and *PaHO/hy1-1*, while they overlapped with Chl autofluorescence in *cHmHO/hy1-1* and *cPaHO/hy1-1*, confirming that *NmHO* and *PaHO* products were localized in the proposed subcellular localizations in each transgenic line.

### Phenotypic analysis of bacterial *HO* transgenic lines

We then performed a phenotypic analysis of the transgenic lines. In 4-day-old seedlings grown under continuous white light, *hy1-1*produced pale-green cotyledons with longer hypocotyls than those of the wild-type Ler ecotype (Fig. 5a). As expected, the expression of BVIXα-producing *NmHO*, either in the cytosol (*NmHO/hy1-1*) or in the plastid (*cNmHO/hy1-1*), complemented the *hy1-1* extended hypocotyl phenotypes (Fig. 5a, b). On the other hand, the introduction of *PaHO* either in cytosol or plastids did not complement the long hypocotyl phenotype of *hy1-1* (Fig. 5a, b), which is consistent with the indication that the functional PHY is not assembled in *PaHO* lines (Fig. 5a, b).

**Figure 5.**
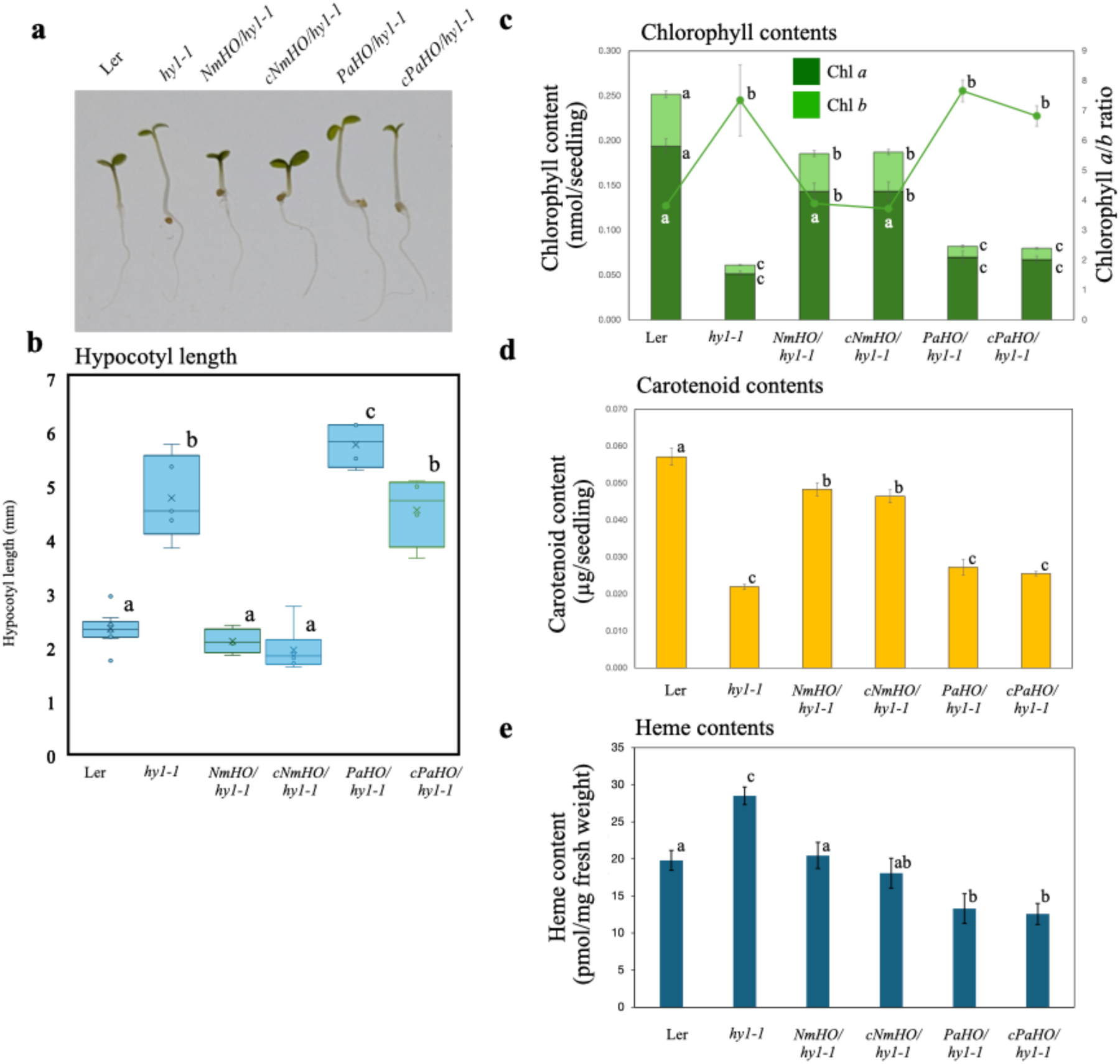
Phenotypic analysis of wild-type Ler, *hy1-1*, and transgenic lines. (a) Photographs of 4-day-old seedlings. (b) Hypocotyl length of each seedling. (c) Chlorophyll contents and chlorophyll *a*/*b* ratio, and (d) total carotenoid contents in transgenic lines. (e) Total heme content. P< 0.05, one-way ANOVA and Tukey’s multiple comparison test.

For pigmentation, *hy1-1* accumulated less Chls and carotenoids compared to those of the wild-type (Fig. 5c, d). Introduction of NmHO either in cytosol or plastids recovered the Chl *a, b*, Chl *a/b* ratio, and carotenoid levels, while those of *PaHO/hy1-1* and *cPaHO/hy1-1* were significantly comparable to those of *hy1-1* (Fig. 5c, d). These results confirm that the introduction of *NmHO* either in plastids or the cytosol can functionally complement the *HO1* deficiency of *hy1-1* by producing BVIXα, leading to functional holo-PHY assembly, whereas BVIXβ/δ-producing PaHO cannot assemble the functional PHY. Meanwhile, the total heme level in *hy1-1* was higher than that in the wild-type Ler (Fig. 5e), consistent with our previous observation in *gun2-1* (Chen et al., 2024). In *NmHO/hy1-1* and *cNmHO/hy1-1*, total heme levels were recovered to the wild-type level. In *PaHO/hy1-1* and *cPaHO/hy1-1*, total heme levels were further decreased compared to the wild-type (Fig. 5e). These results suggest that not only NmHO but also PaHO degrade endogenous heme either in the cytosol or plastids. It is noteworthy that the decreased heme levels were less pronounced than in *Arabidopsis HO1*-introduced *gun2-1* lines, which showed more than 50% reduction (Chen et al., 2024).

### Introduction of *PaHO* restored the *gun* phenotypes

To investigate the impact of regiospecific HO expression on the *gun* phenotype, we examined the transcript levels of representative *PhANGs*: *LIGHT-HARVESTING CHLOROPHYLL-PROTEIN COMPLEX I SUBUNIT A4* (*LHCA4*), *LIGHT-HARVESTING CHLOROPHYLL A/B-BINDING PROTEIN 1.1* (*LHCB1.1*), *LHCB1.2, PHOTOSYSTEM II SUBUNIT QA* (*PSBQA*),

*CARBONIC ANHYDRASE 1* (*CA1*), and *RBCS1A.* With NF treatment, all *PhANGs* expression levels in *hy1-1* were higher than those in Ler (Fig. 6a∼f). In *NmHO/hy1-1* and *cNmHO/hy1-1* lines, the expression levels of all *PhANGs* were significantly suppressed. However, except for *CA1*, the suppressed levels were lower than those of the Ler wild-type, indicating that the *gun* phenotype is partially complemented in these lines. Interestingly, the *gun* phenotype was also complemented in *PaHO/hy1-1* and *cPaHO/hy1-1* lines, showing that heme degradation is essential for complementation of the *gun* phenotype. Notably, in *LHC* genes (Fig. 6a∼c), *cPaHO/hy1-1* showed the most pronounced suppression among the other transgenic lines, suggesting that heme degradation in plastids effectively suppresses *LHC* genes and that this is not related to functional PHY assembly. Suppression levels in remaining genes (*PSBQA*, *CA1*, and *RBCS1A*) in *PaHO/hy1-1* and *cPaHO/hy1-1* were less effective than those in *NmHO/hy1-1* and *cNmHO/hy1-1* lines (Fig. 6d∼f), showing that the mechanism of suppression in these genes is distinct from *LHC* genes.

**Figure 6.**
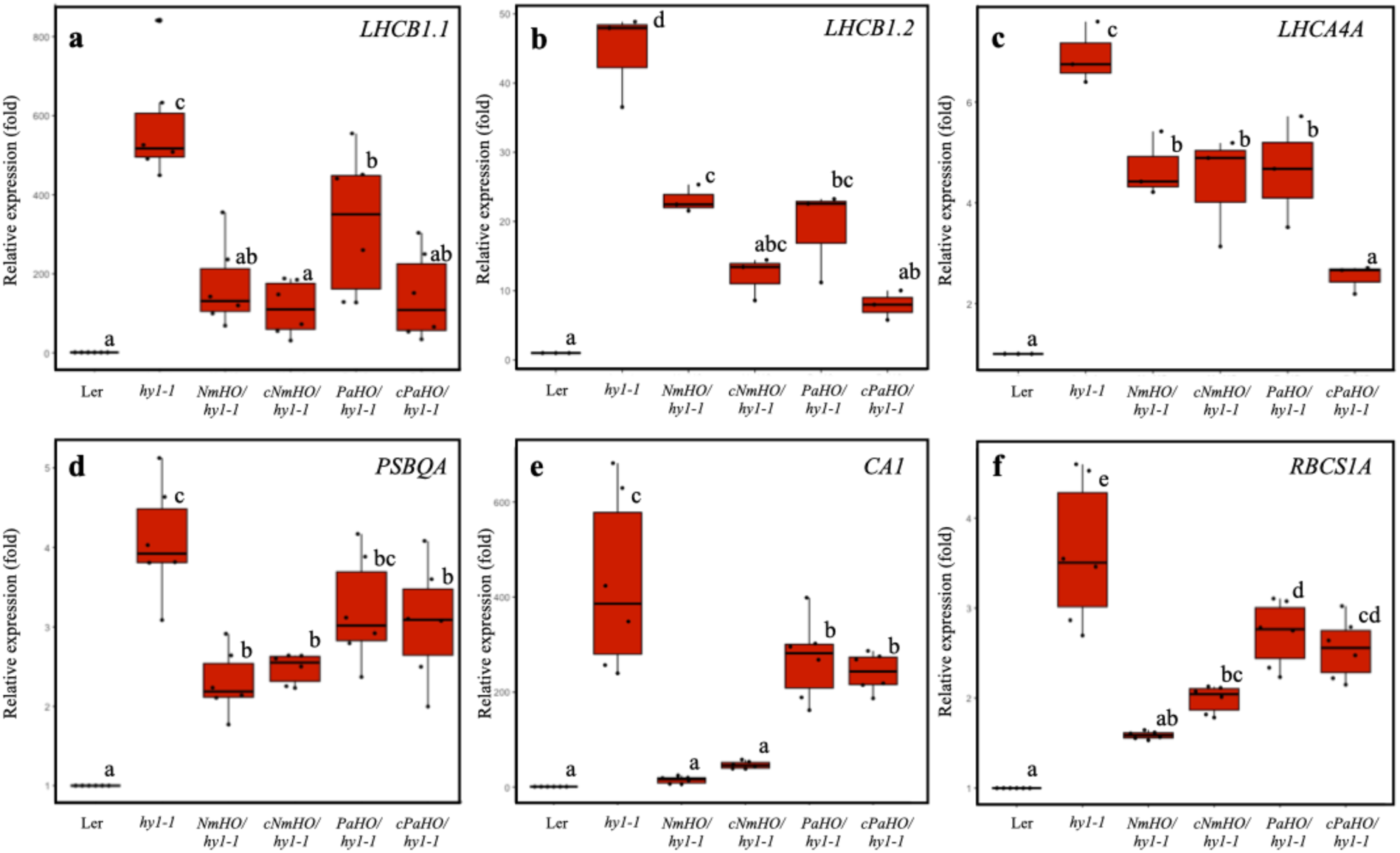
RT-qPCR analysis of 4-day-old seedlings treated with 5 µM norflurazon. Relative transcript levels of (a) *Lhcb1.1*, (b) *Lhcb1.2*, (c) *Lhcb4A*, (d) *PsbQ*, (e) *CA1*, and (f) *RBCS* compared to Ler were depicted. P< 0.05, one-way ANOVA and Tukey’s multiple comparison test.

### Timing of activation of the biogenic retrograde signal

It has been proposed that the biogenic retrograde signals are active during the initial stage of chloroplast development (Pogson et al., 2008). To test this hypothesis, we examined the expression of *FC1, LHCB1.1*, *LHCB1.2, RBCS1A,* and *CA1* genes during 4 days after germination in wild-type Ler (Fig. 7a). Consistent with our previous observation (Espinas et al., 2016), *FC1* is significantly expressed even in 1 day after germination, the level of which is comparable to that of 4 days after germination. After 2 and 3 days of germination, *FC1* expression levels decreased but remained above 20% of the 4-day level. Meanwhile, the expression of *LHCB* genes (*LHCB1.1* and *LHCB1.2*) was activated 3 days after germination (Fig. 7a). Like *FC1*, *RBCS1A* transcripts were detected even in 1 day after germination. After 2 days of germination, *RBCS1A* expression levels decreased but recovered thereafter. Contrastingly, *CA1* transcripts were almost undetectable 1 and 2 days after germination. *CA1* expression was detected thereafter (Fig. 7a). These results show that the regulation mechanisms of expression are distinct among *PhANGs*.

**Figure 7.**
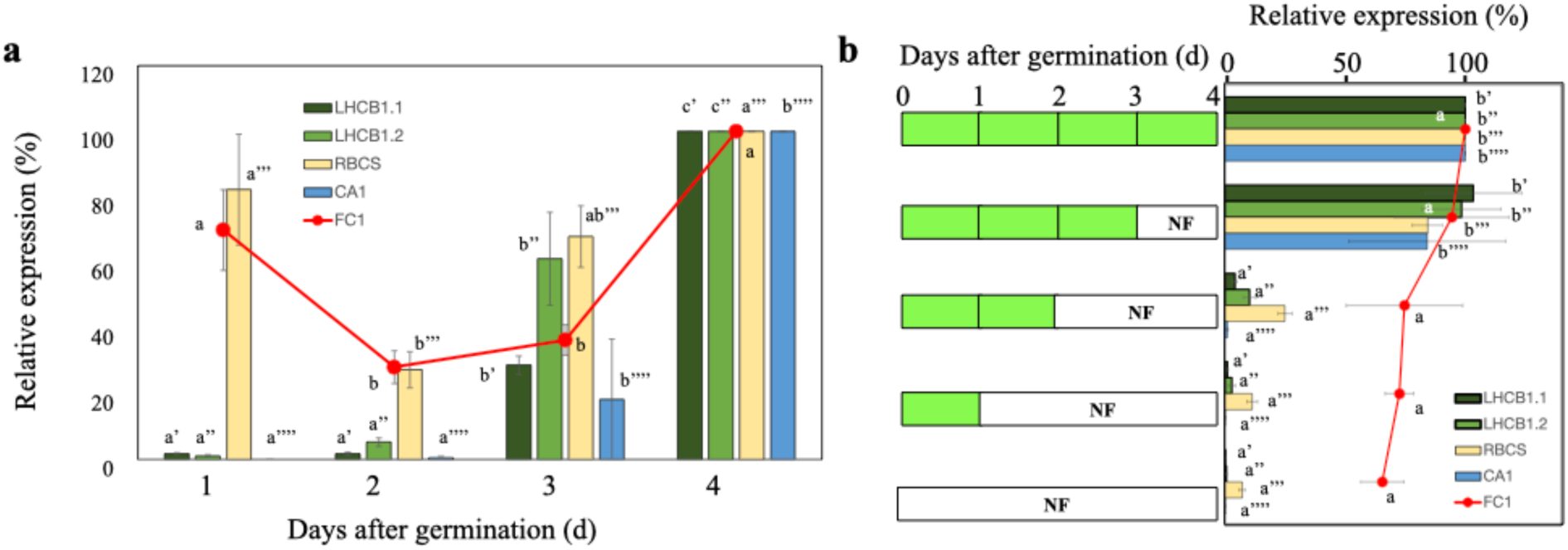
RT-qPCR analysis of *FC1*, *Lhcb1.1, Lhcb1.2, RBCS1A,* and *CA1* during 4 days after germination (a) and 4-day-old seedlings treated with norflurazon (NF) (b, right) with different time regimes (b, left). Relative expression levels of *FC1* (orange, dot), *Lhcb1.1* (dark green, bar), *Lhcb1.2* (green, bar), *RBCS1A* (yellow, bar), and *CA1* (cyan, bar) in the 4-day sample are indicated. P< 0.05, one-way ANOVA and Tukey’s multiple comparison test.

Then, we examined the effect of the addition of NF at different time points after germination (Fig. 7b). When NF was added from the beginning or the end of 1 or 2 days after germination, all *PhANGs* expression was prominently suppressed. There was no noticeable difference in responses among *PhANGs* (Fig. 7b). In comparison, the expression of *FC1* was unaffected (Fig. 7b). However, when NF was added from the end of 3 days after germination, the levels of *PhANGs* expression were the same as those of the untreated control. These results suggest that the FC1-derived heme biogenic signal is activated 3 days after germination.

## Discussion

### Introduction of regiospecific bacterial HOs can dissect the heme and PHY-signaling pathways

To dissect the heme– and PHY-dependent light signaling pathways, we introduced either of the two regiospecific bacterial HOs that produce BVIXα or BVIXβ/δ in *Arabidopsis hy1-1* (Fig. 1). Consistent with previous studies, NmHO and PaHO exhibited regiospecific HO activities *in vitro* (Ratliff et al., 2001). PaHO exhibited higher ascorbate-dependent HO activity than NmHO, as *in vitro* heme degradation proceeded much more rapidly (Fig. 3b, c). HPLC analysis showed that, in contrast to BVIXα-producing NmHO, low levels of BVIXα were detected in PaHO products, while BVIXβ and BVIXδ were produced (Fig. 3e).

Therefore, we hypothesized that the introduction of *PaHO* into *hy1-1* may not assemble functional PHY, since HY2 cannot use BVIXβ/δ as substrate for PΦB biosynthesis. This hypothesis is also supported by the computer-based simulation to estimate the required energy for binding HY2 and BVs (Fig. 2). Based on the docking simulation, we predicted that HY2 preferentially recognizes BVIXα, and that BVIXβ and BVIXδ are unlikely to serve as effective substrates. Consistent with our prediction, in contrast to *NmHO*-introduced lines, *PaHO*-introduced *hy1-1* failed to complement the long hypocotyl and low pigmentation phenotypes (Fig. 5). Since high accumulation of PHYA protein, the level of which is more than 78% of that of *hy1-1*, was detected in *PaHO*-introduced *hy1-1* lines (Fig. 4d), it is evident that functional PHY assembly is almost negligible in these lines.

Consistent with our previous study (Chen et al., 2024), the localization of introduced HO, whether in plastids or the cytosol, was not associated with the basic functionality of bacterial HOs. It is noteworthy that the levels of complementation of *hy1-1* low pigmentation phenotypes by *NmHO* introduction were significantly lower than those observed with wild-type Ler (Fig. 5c, d), contrasting with full complementation of the extended hypocotyl phenotype (Fig. 5a, b). Our previous study showed similar trends: partial complementation of pigmentation but full complementation of hypocotyl length upon introducing *Arabidopsis* cytosolic HO1 (Chen et al., 2024). Therefore, it is likely that these traits have distinct sensitivities to functional PHY assembly. In addition to PHY deficiency, the pigment contents is also affected by feedback inhibition of the heme pool that affects the ALA synthesizing activity (Beale, 1999). Since the wild-type level of heme was detected in *NmHO* lines (Fig. 5e), the differences in pigmentation between wild-type and *NmHO* lines (Fig. 5c, d) is caused by partial PHY assembly as observed higher PHYA protein accumulation in these lines (Fig. 4d).

NmHO introduction into *hy1-1* either the cytosol or plastids recovered the total heme levels to wild-type (Fig. 5e), which is less pronounced than *Arabidopsis* HO1 (Chen et al., 2024), Given that the introduced genes were expressed as proteins in the proposed localizations (Fig. 4c, e), the bacterial NmHO is likely not fully active in *Arabidopsis*, either in plastids or in the cytosol, probably because it cannot utilize sufficient reductant especially plant ferredoxin for the reaction. However, since other traits were entirely (Fig. 5a, b) or partially (Fig. 5c, d) complemented, such lower HO activity can produce functional PHY as observed in ∼75% reduction of PHYA protein compared to *hy1-1* (Fig. 4c). In the *PaHO* lines, much lower levels of total heme were observed (Fig. 5e), consistent with higher *in vitro* HO activity (Fig. 4b, c). This result confirms that heme is actually degraded by *PaHO* introduction. Decreased heme levels in *PaHO* lines (Fig. 5e) were still higher than those in *Arabidopsis* HO1-introduced lines, which showed more than 50% reduction (Chen et al., 2024). By evaluating the above characteristics, we concluded that we can dissect heme signaling and functional PHY assembly using regiospecific bacterial HOs introduced lines.

### Complementation of *gun* phenotype

Introduction of *NmHO* in *hy1-1* complemented the *gun* phenotype (Fig. 6). Except for *CA1*, the levels of *gun* phenotype complementation by *NmHO* introduction were significantly lower than those of wild-type Ler, contrasting with full complementation in *Arabidopsis* HO1 introduction (Chen et al., 2024). However, such partial complementation allowed us to evaluate the involvement of other regulatory signals as discussed below. Interestingly, although the *PaHO* introduction failed to complement the PHY-dependent *hy1-1* phenotypes, it did complement the *hy1-1 gun* phenotype, suggesting that heme degradation is essential for complementation of the *gun* phenotype. Particularly in *LHC* genes (Fig. 6a∼c), the *cPaHO/hy1-1* showed the highest complementation among transgenic lines tested. Furthermore, the complementation levels of *cPaHO/hy1-1* were higher than those of *PaHO/hy1-1*. Since we observed a similar trend with *Arabidopsis HO1* introduction (Chen et al., 2024), it is evident that heme degradation in plastids, the site of heme biosynthesis, is more effective at complementing the *gun* phenotype than in the cytosol.

Since suppression levels in remaining genes (*PSBQA*, *CA1*, and *RBCS1A*) in *PaHO* lines were less effective than those in *NmHO* lines (Fig. 6d∼f), it is assumed that functional PHY assembly is also involved in the *gun* complementation for these *PhANGs*. These results showed that the suppression mechanism differs among the *PhANGs*, which is consistent with a previous report on *Lhcb* and *HEMA1* expression (McCormac and Terry, 2004) and *Lhcb* and *RbcS* expression (Ruckle et al., 2007). In the model presented in the later paper (Ruckle et al., 2007), a complex crosstalk between the repressing retrograde signal and the red light signal pathways has been proposed. Since prominent complementation was observed in *NmHO* lines, functional PHY assembly may suppress *PhANG* expression when the chloroplast is dysfunctional, consistent with the model that red-light photoreceptor(s) suppress *RBCS* expression (Ruckle et al., 2007).

Meanwhile, the FC1-derived heme signal functions as a positive signal for *PhANGs* expression (Woodson et al., 2011; Shimizu and Masuda, 2021). Thus, this model can be updated accordingly.

It is noteworthy that in a green alga, *Chlamydomonas reinhardtii*, the heme catabolites BVIXα and/or phycocyanobilin were found to up-regulate a subset of nuclear genes in darkness, suggesting that bilins function in the light as negative signals to suppress *PhANGs* expression in *C. reinhardtii* (Duanmu et al., 2013). Furthermore, analysis of the *C. reinhardtii HO* mutant (*hmox1*) suggests that plastid bilin biosynthesis is essential for proper regulation of photoacclimation (Wittkopp et al., 2017). Since exogenous BVIXα rescued the *hmox1*’s phenotypic defects in photoautotrophic growth and light-dependent Chl accumulation, it is proposed that a bilin-based retrograde signaling pathway to assemble photosynthetic apparatus is essential for light-growth in *C. reinhardtii*. Here, we showed that *PaHO* lines producing BVIXβ/δ complemented the *gun* phenotype, especially for *LHC* genes. Therefore, it is likely that FC1-derived heme, but not BVIXα, functions as a biogenic retrograde signal, suggesting that a bilin-based retrograde signaling mechanism is not applicable in angiosperms.

### FC1-derived heme functions as a biogenic retrograde signal for initial chloroplast development

In this study, our data confirm that the FC1-derived heme biogenic signal is active during the initial stage of chloroplast development (Pogson et al., 2008). Concerning the initial chloroplast development, it is observed that chloroplast differentiation in cells of the shoot apical meristems occurred 2 day after the onset of germination (Yadav et al., 2019). Using 3-to 4-day-old dark-grown seedlings, the process of chloroplast differentiation during de-etiolation is described (Fujii et al., 2019a; Fujii et al., 2019b; Pipitone et al., 2021). During the de-etiolation process, light activates Chl biosynthesis and mRNA accumulation of *LHCB* genes within 6 h with concomitant increase in galactolipids (Fujii et al., 2019a; Fujii et al., 2019b). Meanwhile, it is proposed that the de-etiolation process can be divided into two phases: the “structure establishment phase (0 ∼24 h)” and the “chloroplast proliferation phase (24 ∼ 96 h)” (Pipitone et al., 2021). The applied light regime (Yadav et al., 2019) or de-etiolation process of etiolated seedlings (Fujii et al., 2019a; Pipitone et al., 2021) is different from ours. If we presume that chloroplast development starts after 2 days (48 h) of germination, the time at which it may correspond to the start of the de-etiolation process, the FC1-derived heme signal becomes active shortly thereafter, during the period of activation of Chl biosynthesis and *LHCB* expression in the structure establishment phase.

It is noteworthy that all *PhANGs* tested showed similar sensitivities to NF treatment (Fig. 7b), despite differing expression profiles (Fig. 7a). Therefore, it is likely that the activations of biogenic retrograde and light signals occur at similar timing.

### Heme trafficking in plant cells

Our data provide clear support for the retrograde heme signaling hypothesis. Currently, the heme trafficking mechanism from organelles is poorly understood in plants compared to yeast and animals (Mochizuki et al., 2010; Shimizu and Masuda, 2021). In yeast, using a genetically encoded fluorescent heme sensor, it was shown that heme synthesized in the inner mitochondrial membrane can be transferred to the nucleus and the cytosol in distinct pathways (Martinez-Guzman et al., 2020). Heme synthesized in the mitochondria was transferred to the nucleus via mitochondria-associated ER membrane contact sites, which was faster than cytosolic heme transfer. Here, our data showed that the plastid-derived heme signal is transferred via the cytosolic pathway to the nucleus in angiosperms. So far, we do not know whether heme for other organelles is transferred via the same pathway. Recently, we demonstrated that the cytosolic cytochrome *b*_5_-like heme-binding protein is required for organ development (Iwata et al., 2025), revealing an unexpected role for heme in plant development. Identification of the components involved in heme trafficking in plants is crucial for a deeper understanding. Furthermore, it is essential to identify the nuclear target of heme.

## Conclusions

By introducing regiospecific bacterial HOs in *Arabidopsis hy1-1*, we demonstrated that heme functions as a retrograde mobile biogenic signal from plastids, passing through the cytosol, to regulate the expression of *PhANG*s, and that this process depends on *PhANGs* for functional PHY assembly. This study significantly advances our understanding of biogenic retrograde signaling in plant cells, which requires initial chloroplast development. Further analysis of the heme trafficking mechanism in plant cells is underway.

## Acknowledgements

We sincerely thank Prof. Tsuyoshi Nakagawa (Interdisciplinary Center for Science Research, Shimane University) for providing the Gateway destination vector pGWB5, Dr. Tsuyoshi Higa (Graduate School of Arts and Sciences, The University of Tokyo) for supporting CLSM observation, and Prof. Matthew J. Terry (School of Biological Sciences, University of Southampton) for critical comments concerning this research.

## Author contributions

M.T. and T.M. designed the research. M.T. performed most of the experiments, and T.S. performed the heme assay. K.M. performed the computer modeling. M.T., K.M., and T.M. analyzed the data. T.M. and K.M. writes the manuscript.

## Supplementary data

**Supplemental Figure 1.** Docking simulation of HY2 with biliverdin isomers under the unrestricted condition

**Supplemental Figure 2.** Codon-optimized sequences of *NmHO* and *PaHO*

**Supplemental Table 1.** List of primers

## Supplementary data

**Supplemental Figure 1.**
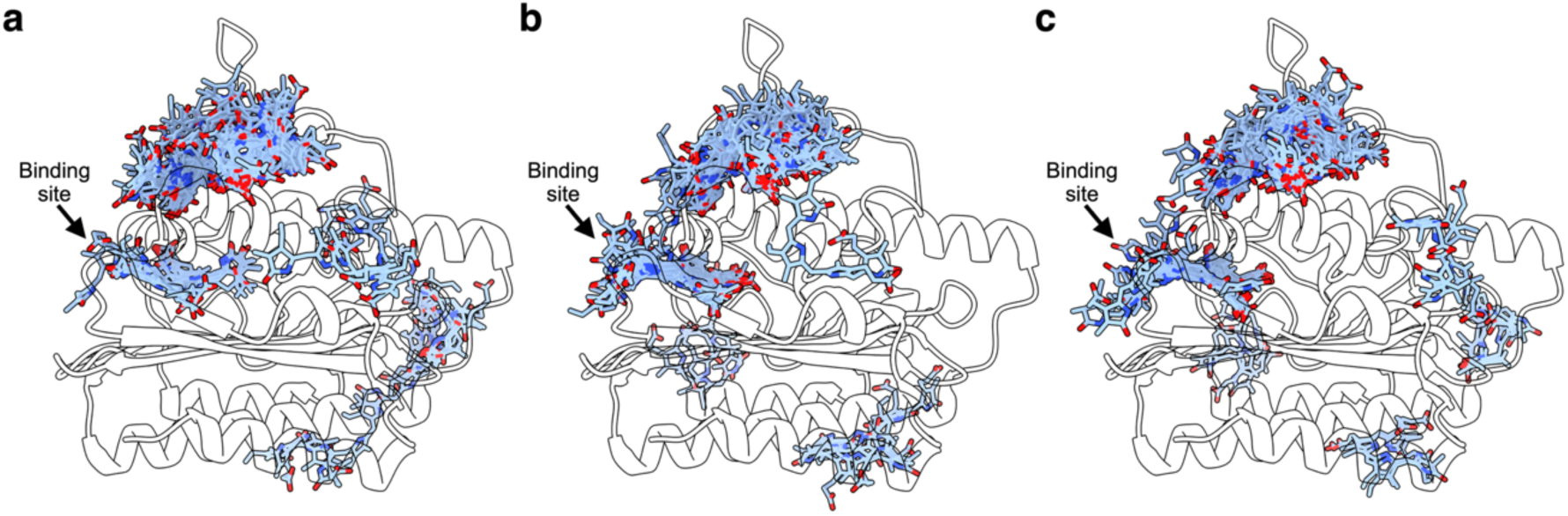
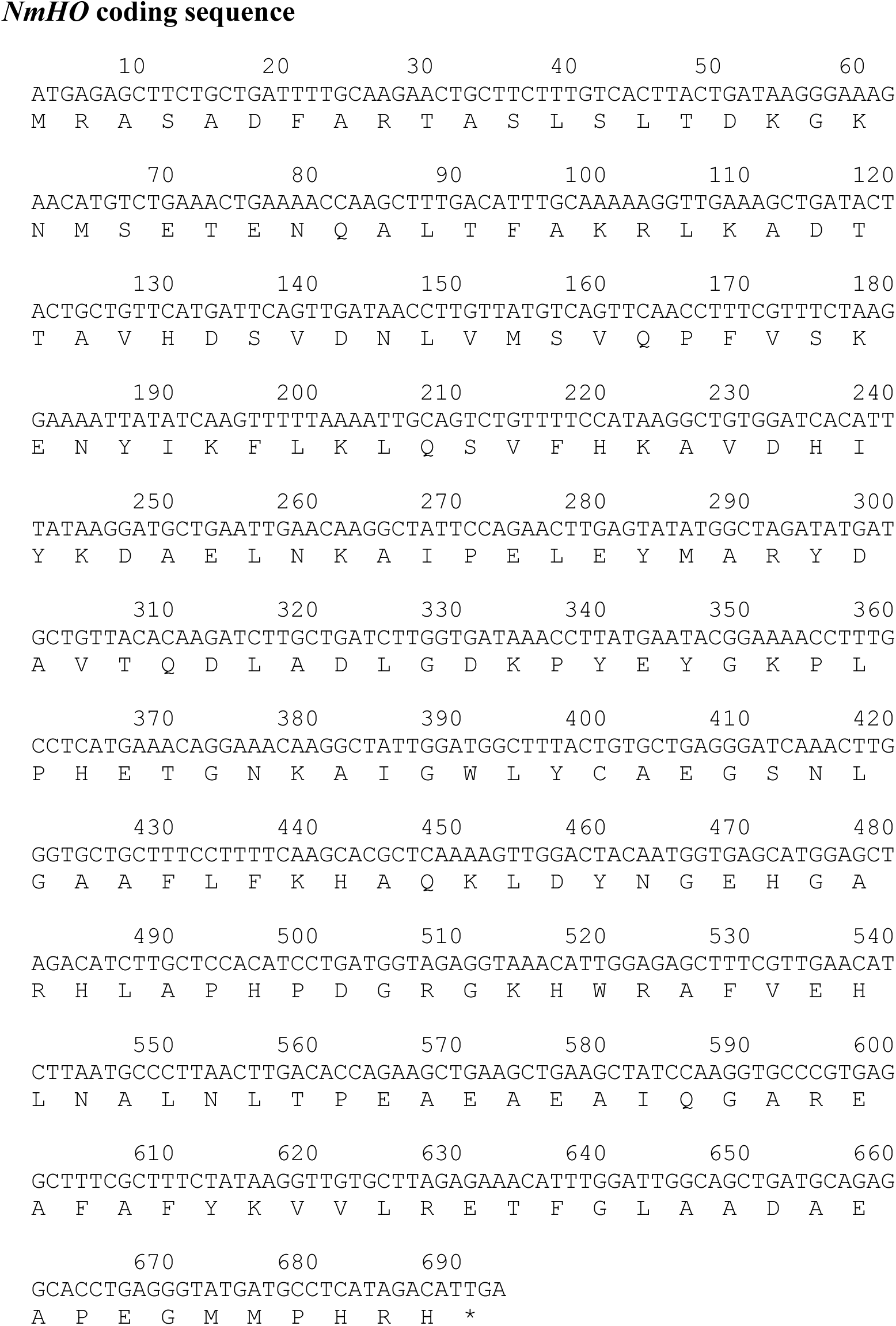
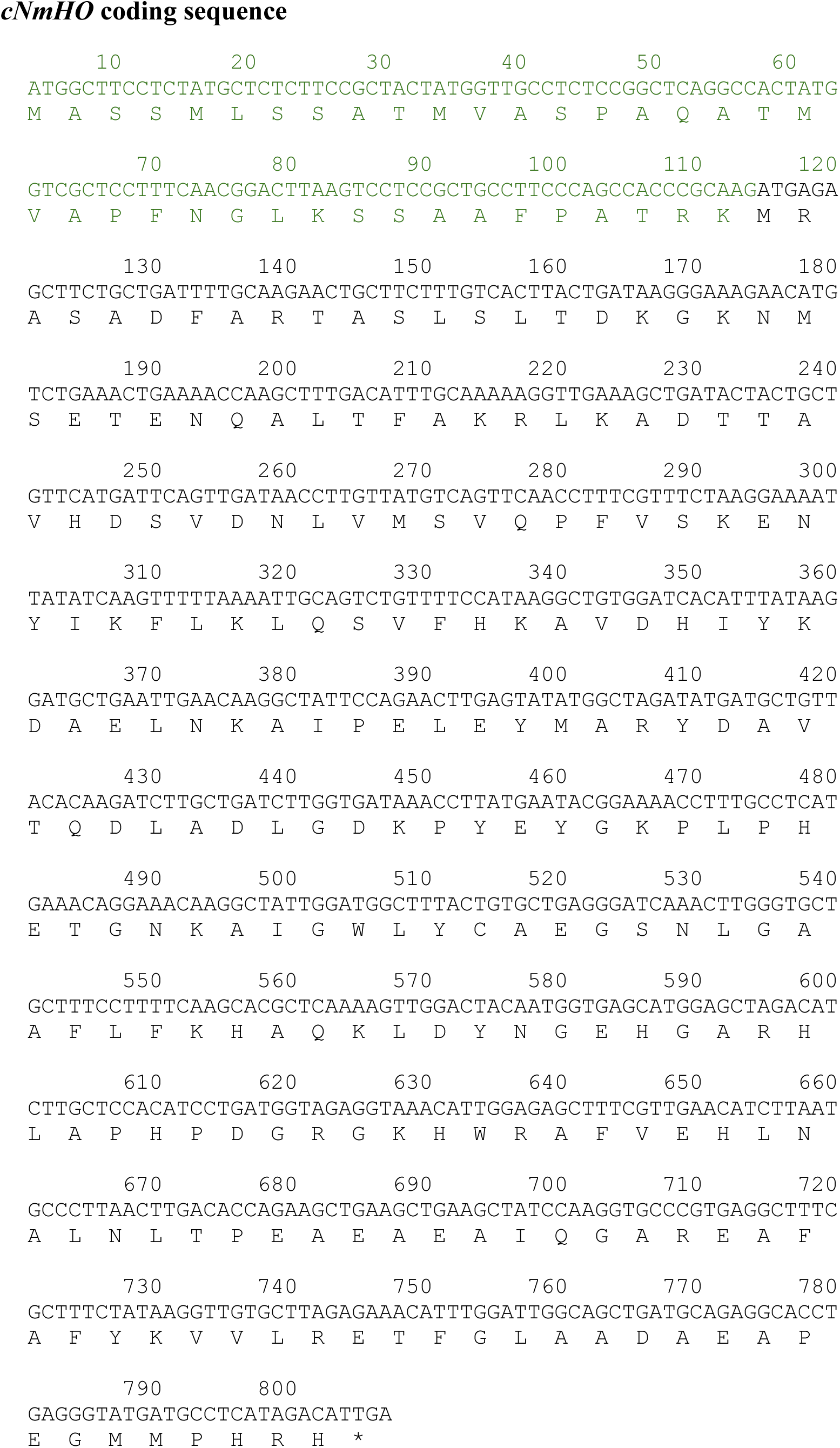
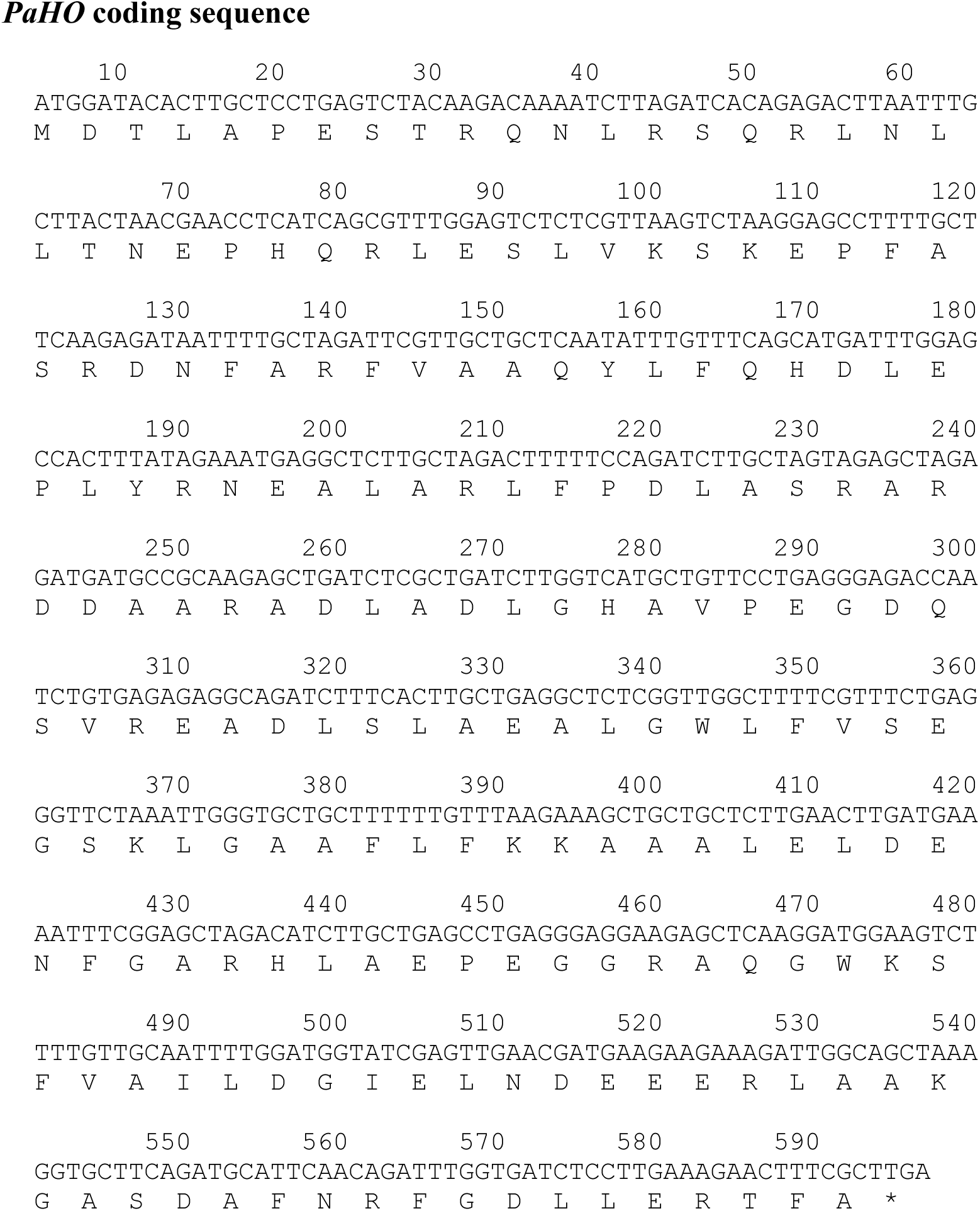
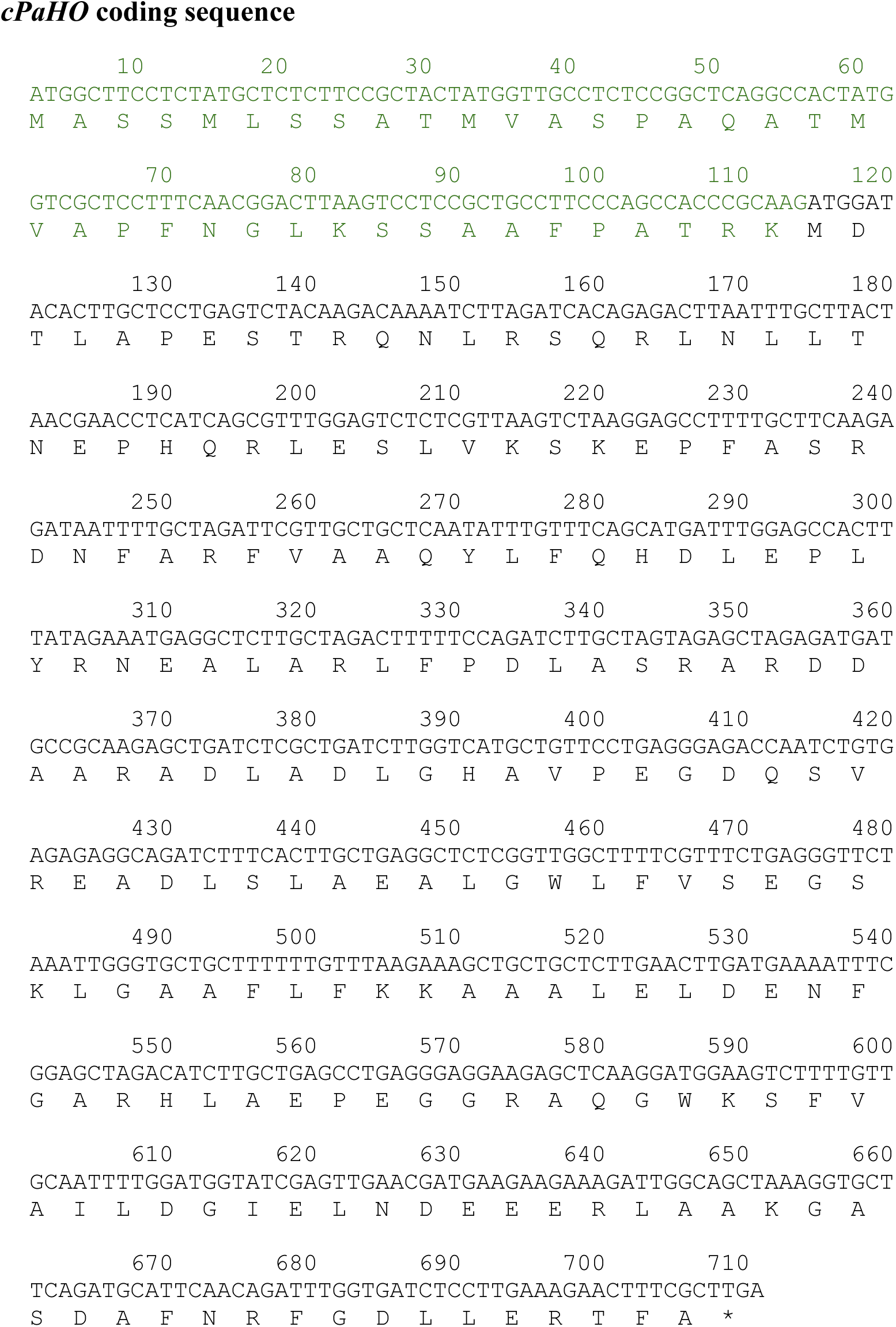
Docking models of Biliverdin Ixα (a), Ixβ (b), and IXδ (c) obtained without spatial restriction (unrestricted docking) in HY2.

**Supplemental** Figure 1 Optimized nucleotide sequences of *NmHO*, *cNmHO*, *PaHO*, and *cPaHO*. Based on the original amino acid sequences (Ratliff et al., 2001), codon-optimized nucleotide sequences for *Arabidopsis* were generated by VectorBuilder (https://en.vectorbuilder.com/). In the cases of cNmHO and cPaHO, the transit peptide sequence of the *Arabidopsis* RBCS (green) was inserted before the initiation codon.

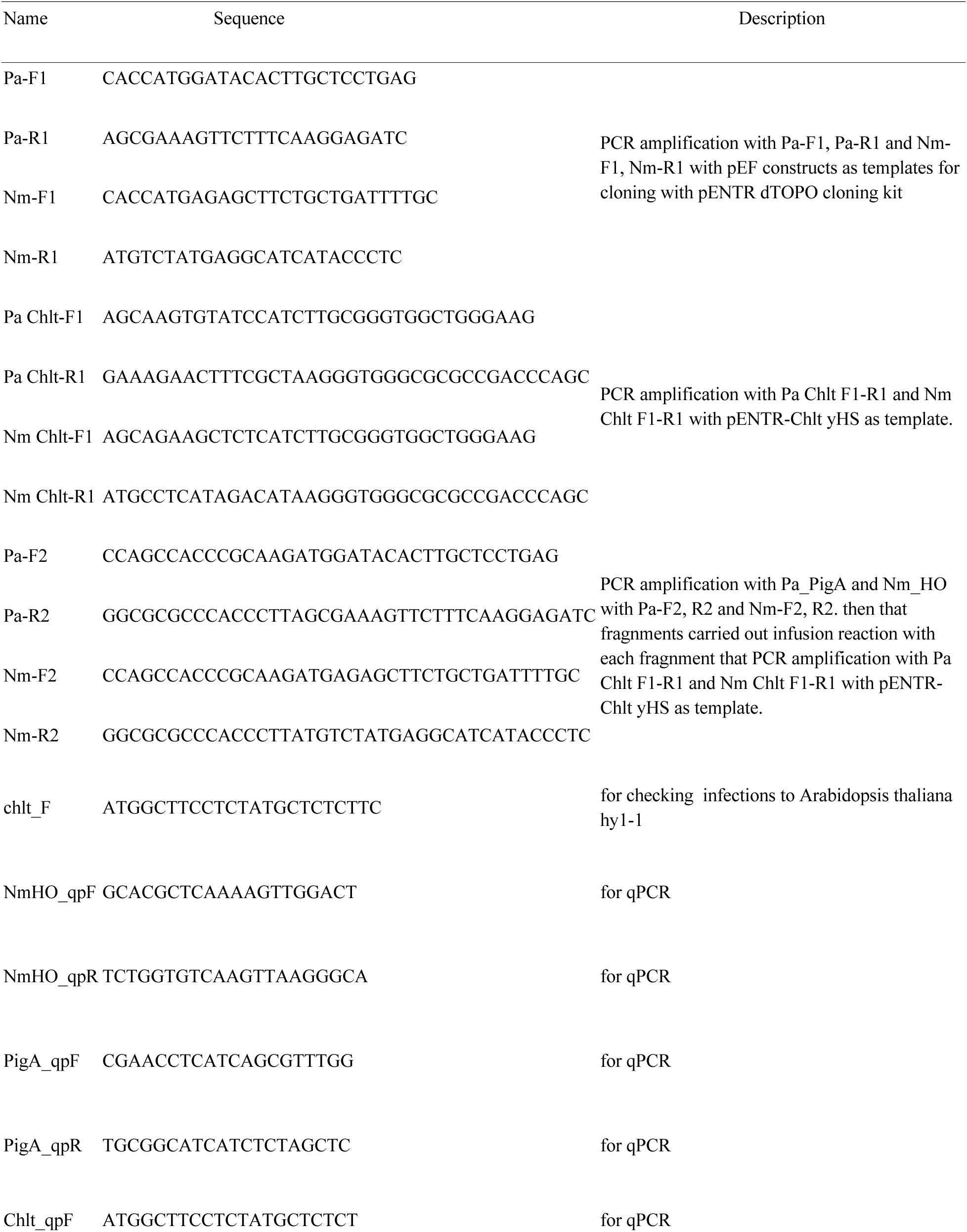
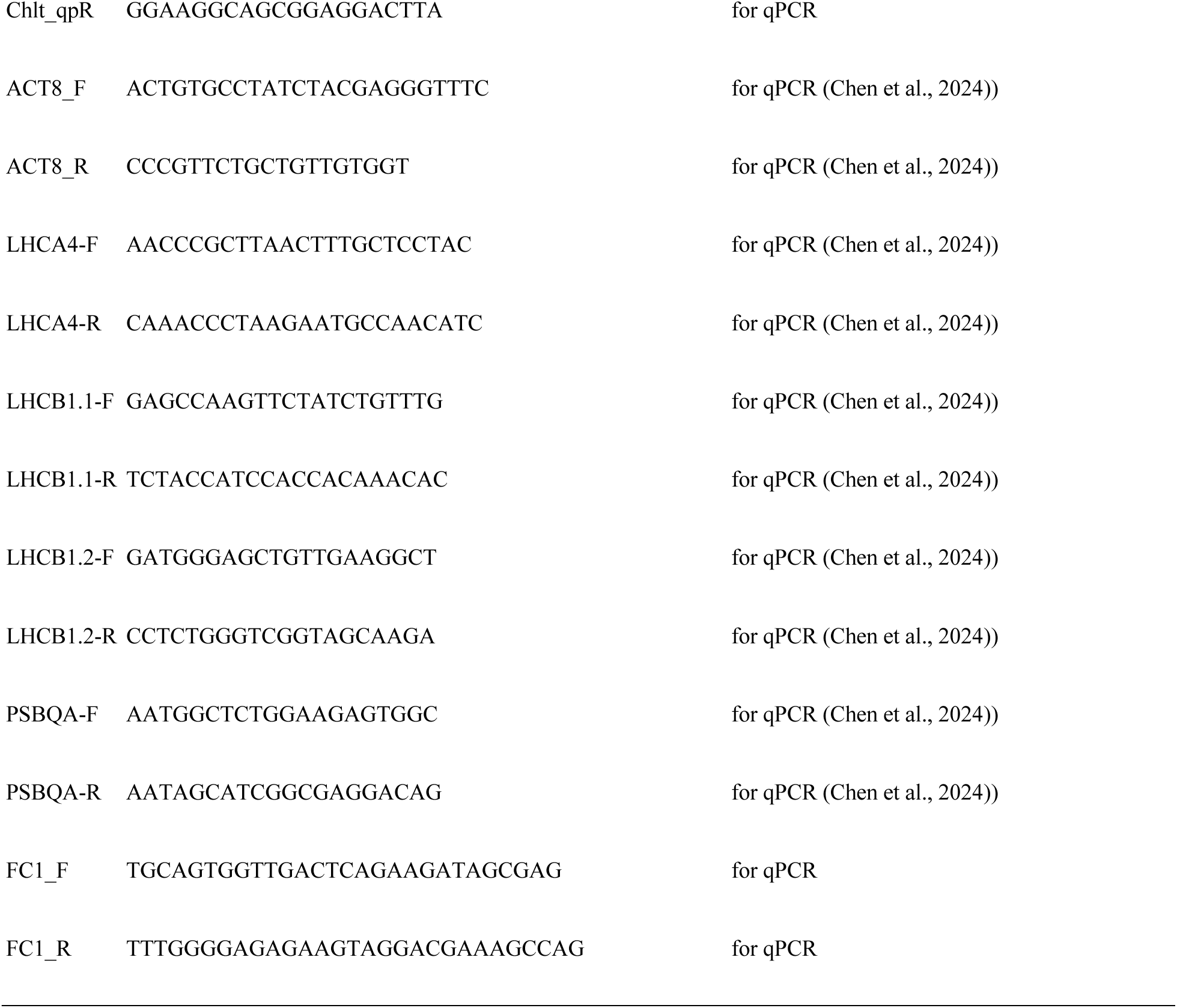
Supplemental Table 1.

