## Supplemental data for "Introduction of regioselective bacterial heme oxygenases into *Arabidopsis hy1-1* supports the retrograde heme signaling hypothesis"

### Supplementary data

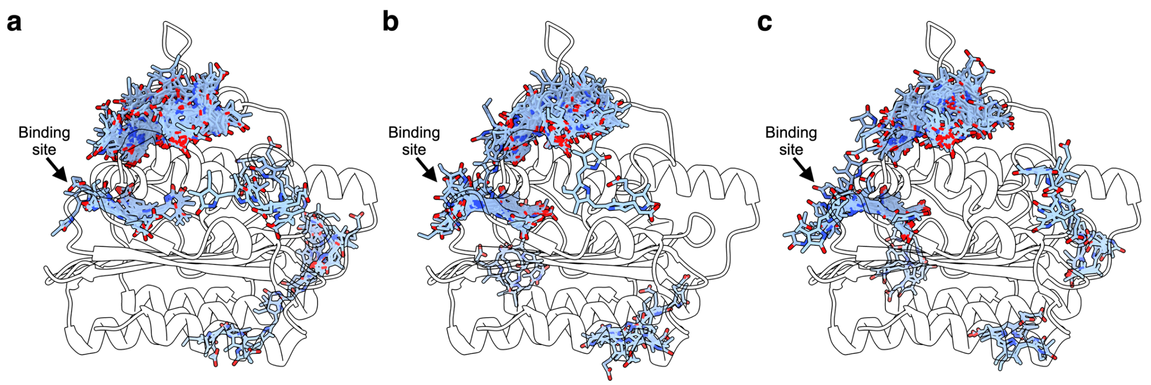


**Supplemental Figure 1**

Docking models of Biliverdin Ixα (a), Ixβ (b), and IXδ (c) obtained without spatial restriction (unrestricted docking) in HY2.

##### *NmHO* coding sequence

```
       10        20        30        40        50        60
ATGAGAGCTTCTGCTGATTTTGCAAGAACTGCTTCTTTGTCACTTACTGATAAGGGAAAG
M  R  A  S  A  D  F  A  R  T  A  S  L  S  L  T  D  K  G  K

         70        80        90       100       110       120
AACATGTCTGAAACTGAAAACCAAGCTTTGACATTTGCAAAAAGGTTGAAAGCTGATACT
N  M  S  E  T  E  N  Q  A  L  T  F  A  K  R  L  K  A  D  T

        130       140       150       160       170       180
ACTGCTGTTCATGATTCAGTTGATAACCTTGTTATGTCAGTTCAACCTTTCGTTTCTAAG
T  A  V  H  D  S  V  D  N  L  V  M  S  V  Q  P  F  V  S  K

        190       200       210       220       230       240
GAAAATTATATCAAGTTTTTAAAATTGCAGTCTGTTTTCCATAAGGCTGTGGATCACATT
E  N  Y  I  K  F  L  K  L  Q  S  V  F  H  K  A  V  D  H  I

        250       260       270       280       290       300
TATAAGGATGCTGAATTGAACAAGGCTATTCCAGAACTTGAGTATATGGCTAGATATGAT
Y  K  D  A  E  L  N  K  A  I  P  E  L  E  Y  M  A  R  Y  D

        310       320       330       340       350       360
GCTGTTACACAAGATCTTGCTGATCTTGGTGATAAACCTTATGAATACGGAAAACCTTTG
A  V  T  Q  D  L  A  D  L  G  D  K  P  Y  E  Y  G  K  P  L

        370       380       390       400       410       420
CCTCATGAAACAGGAAACAAGGCTATTGGATGGCTTTACTGTGCTGAGGGATCAAACTTG
P  H  E  T  G  N  K  A  I  G  W  L  Y  C  A  E  G  S  N  L

        430       440       450       460       470       480
GGTGCTGCTTTCCTTTTCAAGCACGCTCAAAAGTTGGACTACAATGGTGAGCATGGAGCT
G  A  A  F  L  F  K  H  A  Q  K  L  D  Y  N  G  E  H  G  A

        490       500       510       520       530       540
AGACATCTTGCTCCACATCCTGATGGTAGAGGTAAACATTGGAGAGCTTTCGTTGAACAT
R  H  L  A  P  H  P  D  G  R  G  K  H  W  R  A  F  V  E  H

        550       560       570       580       590       600
CTTAATGCCCTTAACTTGACACCAGAAGCTGAAGCTGAAGCTATCCAAGGTGCCCGTGAG
L  N  A  L  N  L  T  P  E  A  E  A  E  A  I  Q  G  A  R  E

        610       620       630       640       650       660
GCTTTCGCTTTCTATAAGGTTGTGCTTAGAGAAACATTTGGATTGGCAGCTGATGCAGAG
A  F  A  F  Y  K  V  V  L  R  E  T  F  G  L  A  A  D  A  E

        670       680       690
GCACCTGAGGGTATGATGCCTCATAGACATTGA
A  P  E  G  M  M  P  H  R  H  *
```

#### *cNmHO* coding sequence

```
        10        20        30        40        50        60
ATGGCTTCCTCTATGCTCTCTTCCGCTACTATGGTTGCCTCTCCGGCTCAGGCCACTATG
M  A  S  S  M  L  S  S  A  T  M  V  A  S  P  A  Q  A  T  M

        70        80        90       100       110       120
GTCGCTCCTTTCAACGGACTTAAGTCCTCCGCTGCCTTCCCAGCCACCCGCAAGATGAGA
V  A  P  F  N  G  L  K  S  S  A  A  F  P  A  T  R  K  M  R

       130       140       150       160       170       180
GCTTCTGCTGATTTTGCAAGAACTGCTTCTTTGTCACTTACTGATAAGGGAAAGAACATG
A  S  A  D  F  A  R  T  A  S  L  S  L  T  D  K  G  K  N  M

       190       200       210       220       230       240
TCTGAAACTGAAAACCAAGCTTTGACATTTGCAAAAAGGTTGAAAGCTGATACTACTGCT
S  E  T  E  N  Q  A  L  T  F  A  K  R  L  K  A  D  T  T  A

       250       260       270       280       290       300
GTTCATGATTCAGTTGATAACCTTGTTATGTCAGTTCAACCTTTCGTTTCTAAGGAAAAT
V  H  D  S  V  D  N  L  V  M  S  V  Q  P  F  V  S  K  E  N

       310       320       330       340       350       360
TATATCAAGTTTTTAAAATTGCAGTCTGTTTTCCATAAGGCTGTGGATCACATTTATAAG
Y  I  K  F  L  K  L  Q  S  V  F  H  K  A  V  D  H  I  Y  K

       370       380       390       400       410       420
GATGCTGAATTGAACAAGGCTATTCCAGAACTTGAGTATATGGCTAGATATGATGCTGTT
D  A  E  L  N  K  A  I  P  E  L  E  Y  M  A  R  Y  D  A  V

       430       440       450       460       470       480
ACACAAGATCTTGCTGATCTTGGTGATAAACCTTATGAATACGGAAAACCTTTGCCTCAT
T  Q  D  L  A  D  L  G  D  K  P  Y  E  Y  G  K  P  L  P  H

       490       500       510       520       530       540
GAAACAGGAAACAAGGCTATTGGATGGCTTTACTGTGCTGAGGGATCAAACTTGGGTGCT
E  T  G  N  K  A  I  G  W  L  Y  C  A  E  G  S  N  L  G  A

       550       560       570       580       590       600
GCTTTCCTTTTCAAGCACGCTCAAAAGTTGGACTACAATGGTGAGCATGGAGCTAGACAT
A  F  L  F  K  H  A  Q  K  L  D  Y  N  G  E  H  G  A  R  H

       610       620       630       640       650       660
CTTGCTCCACATCCTGATGGTAGAGGTAAACATTGGAGAGCTTTCGTTGAACATCTTAAT
L  A  P  H  P  D  G  R  G  K  H  W  R  A  F  V  E  H  L  N

       670       680       690       700       710       720
GCCCTTAACTTGACACCAGAAGCTGAAGCTGAAGCTATCCAAGGTGCCCGTGAGGCTTTC
A  L  N  L  T  P  E  A  E  A  E  A  I  Q  G  A  R  E  A  F

       730       740       750       760       770       780
GCTTTCTATAAGGTTGTGCTTAGAGAAACATTTGGATTGGCAGCTGATGCAGAGGCACCT
A  F  Y  K  V  V  L  R  E  T  F  G  L  A  A  D  A  E  A  P

       790       800
GAGGGTATGATGCCTCATAGACATTGA
E  G  M  M  P  H  R  H  *
```

##### ***PaHO*** **coding sequence**

```
        10         20         30         40         50         60
ATGGATACACTTGCTCCTGAGTCTACAAGACAAAATCTTAGATCACAGAGACTTAATTTG
M  D  T  L  A  P  E  S  T  R  Q  N  L  R  S  Q  R  L  N  L

        70         80         90        100        110        120
CTTACTAACGAACCTCATCAGCGTTTGGAGTCTCTCGTTAAGTCTAAGGAGCCTTTTGCT
L  T  N  E  P  H  Q  R  L  E  S  L  V  K  S  K  E  P  F  A

       130        140        150        160        170        180
TCAAGAGATAATTTTGCTAGATTCGTTGCTGCTCAATATTTGTTTCAGCATGATTTGGAG
S  R  D  N  F  A  R  F  V  A  A  Q  Y  L  F  Q  H  D  L  E

       190        200        210        220        230        240
CCACTTTATAGAAATGAGGCTCTTGCTAGACTTTTTCCAGATCTTGCTAGTAGAGCTAGA
P  L  Y  R  N  E  A  L  A  R  L  F  P  D  L  A  S  R  A  R

       250        260        270        280        290        300
GATGATGCCGCAAGAGCTGATCTCGCTGATCTTGGTCATGCTGTTCCTGAGGGAGACCAA
D  D  A  A  R  A  D  L  A  D  L  G  H  A  V  P  E  G  D  Q

       310        320        330        340        350        360
TCTGTGAGAGAGGCAGATCTTTCACTTGCTGAGGCTCTCGGTTGGCTTTTCGTTTCTGAG
S  V  R  E  A  D  L  S  L  A  E  A  L  G  W  L  F  V  S  E

       370        380        390        400        410        420
GGTTCTAAATTGGGTGCTGCTTTTTTGTTTAAGAAAGCTGCTGCTCTTGAACTTGATGAA
G  S  K  L  G  A  A  F  L  F  K  K  A  A  A  L  E  L  D  E

       430        440        450        460        470        480
AATTTCGGAGCTAGACATCTTGCTGAGCCTGAGGGAGGAAGAGCTCAAGGATGGAAGTCT
N  F  G  A  R  H  L  A  E  P  E  G  G  R  A  Q  G  W  K  S

       490        500        510        520        530        540
TTTGTTGCAATTTTGGATGGTATCGAGTTGAACGATGAAGAAGAAAGATTGGCAGCTAAA
F  V  A  I  L  D  G  I  E  L  N  D  E  E  E  R  L  A  A  K

       550        560        570        580        590
GGTGCTTCAGATGCATTCAACAGATTTGGTGATCTCCTTGAAAGAACTTTCGCTTGA
G  A  S  D  A  F  N  R  F  G  D  L  L  E  R  T  F  A  *
```

##### *cPaHO* coding sequence

```
        10         20         30         40         50         60
ATGGCTTCCTCTATGCTCTCTTCCGCTACTATGGTTGCCTCTCCGGCTCAGGCCACTATG
M  A  S  S  M  L  S  S  A  T  M  V  A  S  P  A  Q  A  T  M

        70         80         90        100        110        120
GTCGCTCCTTTCAACGGACTTAAGTCCTCCGCTGCCTTCCCAGCCACCCGCAAGATGGAT
V  A  P  F  N  G  L  K  S  S  A  A  F  P  A  T  R  K  M  D

       130        140        150        160        170        180
ACACTTGCTCCTGAGTCTACAAGACAAAATCTTAGATCACAGAGACTTAATTTGCTTACT
T  L  A  P  E  S  T  R  Q  N  L  R  S  Q  R  L  N  L  L  T

       190        200        210        220        230        240
AACGAACCTCATCAGCGTTTGGAGTCTCTCGTTAAGTCTAAGGAGCCTTTTGCTTCAAGA
N  E  P  H  Q  R  L  E  S  L  V  K  S  K  E  P  F  A  S  R

       250        260        270        280        290        300
GATAATTTTGCTAGATTCGTTGCTGCTCAATATTTGTTTCAGCATGATTTGGAGCCACTT
D  N  F  A  R  F  V  A  A  Q  Y  L  F  Q  H  D  L  E  P  L

       310        320        330        340        350        360
TATAGAAATGAGGCTCTTGCTAGACTTTTTCCAGATCTTGCTAGTAGAGCTAGAGATGAT
Y  R  N  E  A  L  A  R  L  F  P  D  L  A  S  R  A  R  D  D

       370        380        390        400        410        420
GCCGCAAGAGCTGATCTCGCTGATCTTGGTCATGCTGTTCCTGAGGGAGACCAATCTGTG
A  A  R  A  D  L  A  D  L  G  H  A  V  P  E  G  D  Q  S  V

       430        440        450        460        470        480
AGAGAGGCAGATCTTTCACTTGCTGAGGCTCTCGGTTGGCTTTTCGTTTCTGAGGGTTCT
R  E  A  D  L  S  L  A  E  A  L  G  W  L  F  V  S  E  G  S

       490        500        510        520        530        540
AAATTGGGTGCTGCTTTTTTGTTTAAGAAAGCTGCTGCTCTTGAACTTGATGAAAATTTC
K  L  G  A  A  F  L  F  K  K  A  A  A  L  E  L  D  E  N  F

       550        560        570        580        590        600
GGAGCTAGACATCTTGCTGAGCCTGAGGGAGGAAGAGCTCAAGGATGGAAGTCTTTTGTT
G  A  R  H  L  A  E  P  E  G  G  R  A  Q  G  W  K  S  F  V

       610        620        630        640        650        660
GCAATTTTGGATGGTATCGAGTTGAACGATGAAGAAGAAAGATTGGCAGCTAAAGGTGCT
A  I  L  D  G  I  E  L  N  D  E  E  E  R  L  A  A  K  G  A

       670        680        690        700        710
TCAGATGCATTCAACAGATTTGGTGATCTCCTTGAAAGAACTTTCGCTTGA
S  D  A  F  N  R  F  G  D  L  L  E  R  T  F  A  *
```

**Supplemental Figure 1**

Optimized nucleotide sequences of *NmHO*, *cNmHO*, *PaHO*, and *cPaHO*. Based on the original amino acid sequences (Ratliff et al., 2001), codon-optimized nucleotide sequences for *Arabidopsis* were generated by VectorBuilder (https://en.vectorbuilder.com/). In the cases of cNmHO and cPaHO, the transit peptide sequence of the *Arabidopsis* RBCS (green) was inserted before the initiation codon.

#### Supplemental Table 1

| Name | Sequence | Description |
|---|---|---|
| Pa-F1 | CACCATGGATACACTTGCTCCTGAG | |
| Pa-R1 | AGCGAAAGTTCTTTCAAGGAGATC | PCR amplification with Pa-F1, Pa-R1 and Nm-F1, Nm-R1 with pEF constructs as templates for cloning with pENTR dTOPO cloning kit |
| Nm-F1 | CACCATGAGAGCTTCTGCTGATTTTGC | |
| Nm-R1 | ATGTCTATGAGGCATCATACCCTC | |
| Pa Chlt-F1 | AGCAAGTGTATCCATCTTGCGGGTGGCTGGGAAG | |
| Pa Chlt-R1 | GAAAGAACTTTCGCTAAGGGTGGGCGCGCCGACCCAGC | PCR amplification with Pa Chlt F1-R1 and Nm Chlt F1-R1 with pENTR-Chlt yHS as template. |
| Nm Chlt-F1 | AGCAGAAGCTCTCATCTTGCGGGTGGCTGGGAAG | |
| Nm Chlt-R1 | ATGCCTCATAGACATAAGGGTGGGCGCGCCGACCCAGC | |
| Pa-F2 | CCAGCCACCCGCAAGATGGATACACTTGCTCCTGAG | |
| Pa-R2 | GGCGCGCCCACCCTTAGCGAAAGTTCTTTCAAGGAGATC | PCR amplification with Pa_PigA and Nm_HO with Pa-F2, R2 and Nm-F2, R2. then that fragnments carried out infusion reaction with each fragnment that PCR amplification with Pa Chlt F1-R1 and Nm Chlt F1-R1 with pENTR-Chlt yHS as template. |
| Nm-F2 | CCAGCCACCCGCAAGATGAGAGCTTCTGCTGATTTTGC | |
| Nm-R2 | GGCGCGCCCACCCTTATGTCTATGAGGCATCATACCCTC | |
| chlt_F | ATGGCTTCCTCTATGCTCTCTTC | for checking infections to Arabidopsis thaliana hy1-1 |
| NmHO_qpF | GCACGCTCAAAAGTTGGACT | for qPCR |
| NmHO_qpR | TCTGGTGTCAAGTTAAGGGCA | for qPCR |
| PigA_qpF | CGAACCTCATCAGCGTTTGG | for qPCR |
| PigA_qpR | TGCGGCATCATCTCTAGCTC | for qPCR |
| Chlt_qpF | ATGGCTTCCTCTATGCTCTCT | for qPCR |

| | | |
|---|---|---|
| Chlt_qpR | GGAAGGCAGCGGAGGACTTA | for qPCR |
| ACT8_F | ACTGTGCCTATCTACGAGGGTTTC | for qPCR (Chen et al., 2024)) |
| ACT8_R | CCCGTTCTGCTGTTGTGGT | for qPCR (Chen et al., 2024)) |
| LHCA4-F | AACCCGCTTAACTTTGCTCCTAC | for qPCR (Chen et al., 2024)) |
| LHCA4-R | CAAACCCTAAGAATGCCAACATC | for qPCR (Chen et al., 2024)) |
| LHCB1.1-F | GAGCCAAGTTCTATCTGTTTG | for qPCR (Chen et al., 2024)) |
| LHCB1.1-R | TCTACCATCCACCACAAACAC | for qPCR (Chen et al., 2024)) |
| LHCB1.2-F | GATGGGAGCTGTTGAAGGCT | for qPCR (Chen et al., 2024)) |
| LHCB1.2-R | CCTCTGGGTCGGTAGCAAGA | for qPCR (Chen et al., 2024)) |
| PSBQA-F | AATGGCTCTGGAAGAGTGGC | for qPCR (Chen et al., 2024)) |
| PSBQA-R | AATAGCATCGGCGAGGACAG | for qPCR (Chen et al., 2024)) |
| FC1_F | TGCAGTGGTTGACTCAGAAGATAGCGAG | for qPCR |
| FC1_R | TTTGGGGAGAGAAGTAGGACGAAAGCCAG | for qPCR |
